# Asymmetrical recognition and processing of double-strand breaks formed during DNA replication

**DOI:** 10.1101/2025.07.03.663015

**Authors:** Matthew J Johnson, Michael T Kimble, Seoyeon Jeong, Aakanksha Sane, Lorraine S Symington

**Author notes:** Corresponding author: Lorraine S Symington. **Author contributions:** Conceptualization, M.J.J. and L.S.S.; Methodology, M.J.J., M.T.K., S.Y.J, A.S., and L.S.S.; Formal Analysis, M.J.J., M.T.K., S.Y.J, A.S., and L.S.S.; Investigation, M.J.J., M.T.K., S.Y.J., A.S.; Writing, M.J.J. and L.S.S.; Visualization, M.J.J. and L.S.S.; Supervision, L.S.S.; Funding Acquisition, L.S.S. **Competing interests statement:** The authors declare no competing interests.

## Abstract

DNA end resection to generate 3′ ssDNA overhangs is the first step in homology-directed mechanisms of double-strand break (DSB) repair. While end resection has been extensively studied in the repair of endonuclease-induced DSBs, little is known about how resection proceeds at DSBs generated during DNA replication. We previously established a system to generate replication-dependent double-ended DSBs at the sites of nicks induced by the Cas9^D10A^ nickase in the budding yeast genome. Here, we suggest that these DSBs form in an asymmetric manner, with one break end being blunt or near blunt, and the other bearing a 3′ ssDNA overhang of up the size of an Okazaki fragment. We find that Mre11 preferentially binds blunt ends and is required for the removal Ku from these DSB ends. In contrast, the ends predicted to have 3′ overhangs have minimal Ku binding, and end resection at these break ends can proceed in a mostly Mre11-independent manner through either the Exo1 or Dna2-Sgs1 long-range resection pathways. These findings indicate that resection proceeds differently at replication-dependent DSBs than at canonical DSBs, and reveals that Ku selectively binds nearly blunt ends, potentially explaining why replication-dependent DSBs are poorly repaired by non-homologous end joining.

**Significance Statement:** DNA end resection is a critical early step in the repair of DNA double-strand breaks by homologous recombination. Studies using endonucleases that generate DSBs in all phases of the cell cycle have shown that end resection proceeds by the sequential action of the Mre11 complex and long-range resection nucleases, Exo1 or Dna2-Sgs1. In this study, we restricted DSB generation to S-phase and found that the genetic requirements for end resection at more physiologically relevant DSBs are different to those at endonuclease-induced DSBs. The asymmetrical recognition and processing of replication-dependent DSBs may also explain the bias toward homologous recombination mediated repair over non-homologous end joining.

## Introduction

DNA double strand breaks (DSBs) are cytotoxic DNA lesions that must be repaired to maintain genome integrity. DSBs can be generated directly using site-specific endonucleases, such as I-SceI, HO, and AsiSI (1, 2), or produced randomly in the genomic, for example, by ionizing radiation or use of drugs that trap Topoisomerase II cleavage complexes (3, 4). Studies using endonucleases have provided a general framework for how DSBs are recognized and repaired (5, 6). These studies suggest that cells initially attempt repair by non-homologous end joining (NHEJ) but if the DSB ends are not compatible for ligation, and cells are in the appropriate cell cycle phase, they undergo end resection to repair by homologous recombination (HR) (7–9). End resection is initiated by the Mre11-Rad50-Xrs2/Nbs1 (MRX in *S. cerevisiae*, MRN in other eukaryotes) complex with its cofactor, phosphorylated Sae2, to form short 3′ overhangs (10). Overhangs generated by MRX are then extended by the long-range resection machinery, including Exo1 and the Dna2-Sgs1-Top3-Rmi1 ensemble (referred to as Dna2-Sgs1 hereafter) (10). The resulting single-stranded DNA (ssDNA) is initially bound by RPA and then replaced by Rad51 to promote strand exchange with an intact donor sequence, followed by repair synthesis to replace sequences lost by end resection (9, 11).

While studies with site-specific endonucleases have been invaluable in deciphering DSB repair pathways (5), it is increasingly recognized that endonuclease-induced DSBs do not accurately mimic DNA damage generated during DNA replication. Given that replication-dependent DNA damage is predicted to be quite common and a driver of genome instability (12, 13), further research is needed to fully understand DNA repair processes in the context of DNA replication. Single-stranded breaks (SSBs), also called nicks, are common intermediates in DNA metabolic processes and have the potential to be converted to DSBs by replication fork progression (12, 14). Indeed, some cancer therapeutics act by inducing or stabilizing SSBs resulting in the formation of cytotoxic DSBs during S phase (15–17). An *in vitro* study using Xenopus egg extracts and pre-nicked plasmid templates demonstrated formation of seDSBs by replisome run off at nicks, regardless of whether the nick was on the leading or lagging-strand template. However, the end structure of the seDSB differed for leading and lagging strand collapses. Replisome run off at a nick on the leading-strand template generated an end with a 3 nucleotide (nt) 5′ overhang, referred to here as a blunt end, whereas the seDSB produced by a lagging strand collapse had a 3′ overhang of ∼70 nt.

Recent work has demonstrated the utility of modified CRISPR/Cas9 enzymes for generation of strand specific nicks that can be converted to DSBs during replication (18–23). While Cas9 makes a DSB through cooperative action of two endonuclease domains, Cas9^D10A^ harbors a mutation in one of these two domains (24) rendering the enzyme able to cut only one strand the DNA duplex. In human cells, nicks generated by Cas9^D10A^ on the leading-strand template were shown to convert to mostly seDSBs during S phase, while nicks on the lagging-strand template resulted in double-ended DSBs (deDSBs) (18). Formation of a deDSB was shown to result from continued translocation of the CMG replicative helicase on the leading strand, bypassing the nick on the lagging-strand template (18). An Okazaki fragment initiated after replisome bypass would be predicted to terminate at the nick, yielding a deDSB (Fig S1A). In budding and fission yeasts, nicks formed by Cas9^D10A^, M13 gpII nickase or a minimal HO cut site that is preferentially nicked, generate mostly deDSBs during replication, even for nicks generated on the leading strand (19, 20, 25). In the case of the Flp-nick system (26), there are conflicting reports on whether single end or deDSBs are formed (20, 27–29). The deDSB signal detected for leading-strand template nicks is thought to be due to convergence of two opposing forks at the nick. Such a scenario might be more likely in organisms such as yeasts with a high density of active and dormant origins (30).

Genetic analysis in *Saccharomyces cerevisiae* revealed that Cas9^D10A^-induced replication-coupled DSBs are repaired by Rad51-dependent sister-chromatid recombination with no apparent contribution by NHEJ (19). This system therefore provides the opportunity to study sister-chromatid recombination in a replication context, potentially revealing mechanistic aspects of HR-mediated repair that have gone undetected in studies of canonical DSB repair using endonucleases such as HO or I-SceI. In this study, we address how the ends of replication-dependent DSBs are recognized and repaired. Since work using the Xenopus egg extract system showed different end structures for seDSBs produced at nicks on leading and lagging strand templates (31), we investigated whether the ends of a replication-dependent deDSB are asymmetrical, consisting of one blunt end and the other with a 3′ overhang. We find that resection at blunt ends is dependent on MRX mediated short-range resection. In contrast, resection at 3′ overhang ends can proceed in a mostly MRX independent manner by either Exo1 or Dna2-Sgs1-catalyzed long-range resection. We also find that the Ku complex is recruited to blunt ends of replication dependent DSBs, and that the removal of the Ku complex by MRX is required for repair of breaks. Taken together, these results paint a more nuanced picture of resection dynamics at breaks formed during replication and highlight the significance of the structural asymmetry of these breaks.

## Results

### MRX nuclease activity is dispensable for repair of replication-dependent DSBs

To study breaks derived from Cas9^D10A^ initiated nicks, we used a set of four guides flanking *ARS607*, a highly efficient, early-firing origin on chromosome VI of *S. cerevisiae* (Figure 1A) (19). Guide RNAs (gRNA) 1 and 20 generate nicks on the lagging-strand template, while gRNAs 6 and 14 generate nicks on the leading-strand template when replication originates from *ARS607* (Figure S1A) (19). In previous work we found that targeting the nickase with gRNAs 1, 6, and 20 results in replication-dependent deDSBs, while targeting the nickase with gRNA14 generates a mostly seDSB signal (Figure S1A) (19). Since HR-deficient mutants were previously reported to show the highest sensitivity to gRNA1 and gRNA6 (19), we focused on these gRNAs for most assays in this study. We first verified that each member of the MRX complex is required for cell viability to nicks produced on leading and lagging strand templates (Figure 1B, 1C, S1B). In contrast, the *mre11-H125N* allele, in which the MRX complex lacks nuclease activity (32), conveyed only mild sensitivity to gRNA1 and no sensitivity to gRNA6 (Figure 1C). We also tested the involvement of Sae2 and Tel1, which stimulate MRX resection initiation and signal the DNA damage checkpoint via MRX (33), respectively (Figure 1B). However, neither of these mutants displayed detectable nickase sensitivity (Figure 1C). These results demonstrate that MRX nuclease activity is not *per se* required for replication-dependent DSB repair, in agreement with past work in *S. cerevisiae* studying repair of DSB ends lacking covalently bound proteins (32).

**Figure 1.**
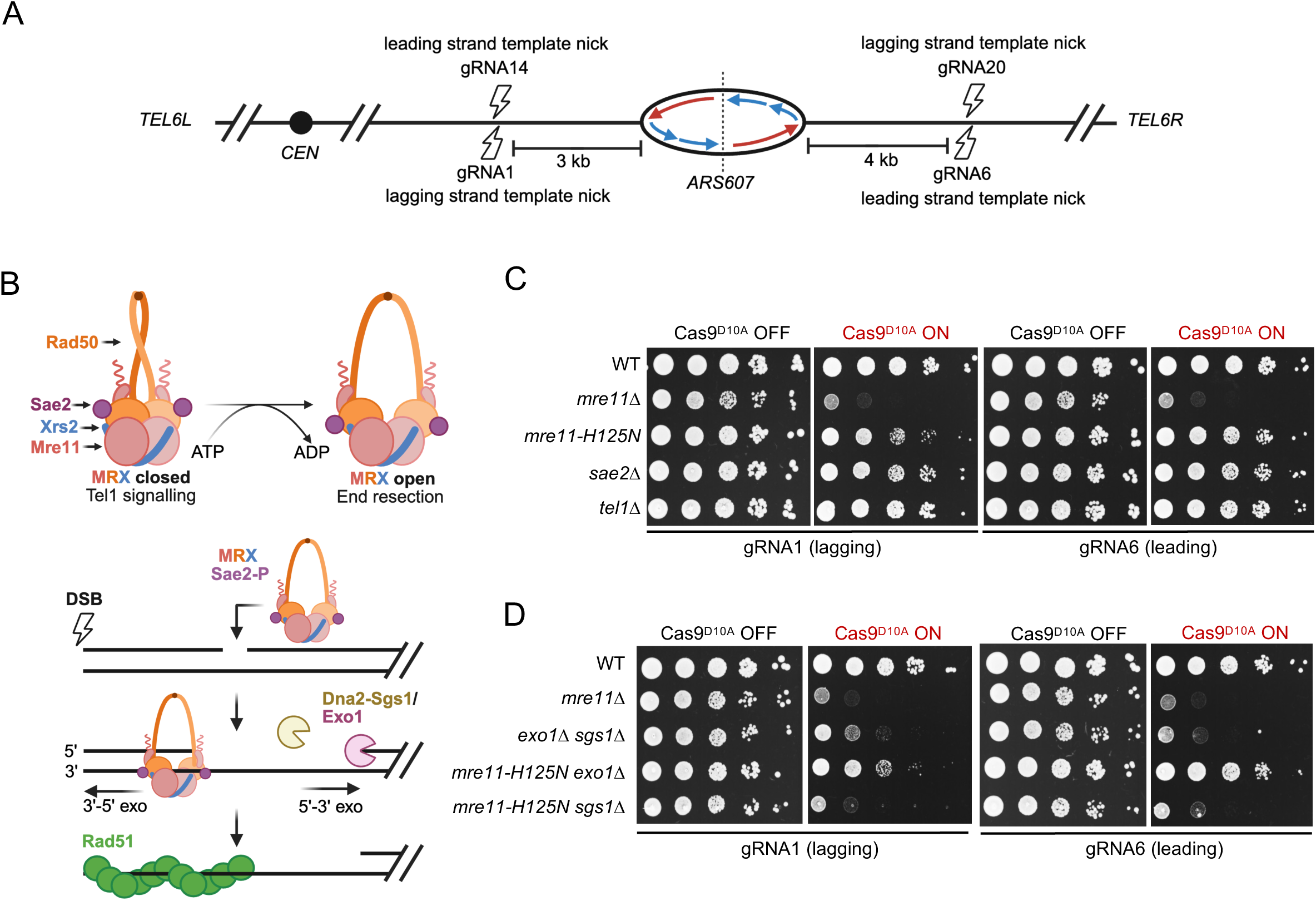
MRX nuclease activity is not required for replication-dependent DSB repair. **A.** Chromosomal location of the main gRNAs used to generate replication-dependent DSBs in this study. gRNA1 and gRNA20 generate nicks on the lagging-strand template, while gRNA6 and gRNA14 generate leading-strand template nicks with Cas9^D10A^. **B.** Schematic of the structure and roles of the MRX complex highlighting the ability of the complex to initiate end resection in a Sae2 dependent manner and to promote Tel1 signaling (top). Schematic for DNA end resection showing one side of a DSB (indicated by a lightning bolt). MRX, stimulated by phosphorylated Sae2 incises the 5’-terminated strand near a DSB to create the entry site for bi-directional resection using the Mre11 3’-5’ exonuclease activity and either Exo1 or Dna2-Sgs1 nucleases (bottom). **C., D.** Ten-fold **s**erial dilutions of the indicated strains expressing gRNA1 or gRNA6 were spotted on medium -/+ β-estradiol to induce Cas9^D10A^ expression.

To further define the mechanism for resection of replication-dependent DSBs, we generated double mutants with *mre11-H125N* and either *exo1*Δ or *sgs1*Δ. Because the MRX complex is known to physically recruit Dna2-Sgs1 to DSBs in addition to facilitating short-range resection (34–36), we reasoned that the *mre11-H125N sgs1*Δ mutant would be highly sensitive to nickase induction. We found this to be the case for breaks generated by gRNAs 1, 6, 14 and 20 (Figure 1D and Figure S1C), while the *mre11-H125N exo1*Δ mutant showed sensitivity comparable to the *mre11-H125N* single mutant (Figure 1D). Curiously, the *mre11-H125N* and *mre11-H125N exo1Δ* mutants both seem to be slightly more sensitive to nicks on the lagging strand rather than the leading-strand template (Figure 1C, 1D, and Figure S1C). These genetic assays support a previous study reporting the requirement for Dna2-Sgs1 dependent resection in the absence of MRX nuclease activity for repair of ionizing radiation induced DSBs (37). Additionally, we verified that the single deletions of *exo1Δ* or *sgs1Δ* do not confer nickase sensitivity whereas loss of long-range end resection renders cells sensitive to nickase induction (Figure 1D, and Figure S1B, C), as previously reported (19, 38, 39).

### Replication-dependent DSBs are likely asymmetrical

We next wished to determine whether the structure of replication-dependent DSBs is changed in the absence of Mre11. We expressed Cas9^D10A^ with gRNA6 or gRNA20 in asynchronous populations of WT or *mre11Δ* cells and analyzed genomic DNA by restriction endonuclease digestion followed by Southern blot analysis (Figure 2A). As reported previously, these gRNAs generate a mostly deDSB signal in WT cells (19). Surprisingly, *mre11Δ* cells appear to generate seDSBs for both targeted sites when cut with Cas9^D10A^, but with the opposite polarity (Figure 2B). Given that the Cas9^D10A^ nicks introduced with these gRNAs are only 7 nt apart on opposite strands, they should both be replicated by a unidirectional replication fork initiated from *ARS607*. For both gRNAs, the end that persists in *mre11Δ* cells corresponds to the one predicted to be blunt. This same trend of a predominantly seDSB signal but with opposite polarity was observed when targeting Cas9^D10A^ with gRNA1 and gRNA1 with in *mre11Δ* cells (Figure S2A, B). We suggest that the observed pattern of one persistent break fragment results from resection of the other DSB end, which would prevent restriction enzyme cleavage and lead to fragment disappearance. Since the role of MRX is to generate 3′ overhang ends for Exo1 and/or Dna2-Sgs1 to further resect, the most plausible explanation for this result is that resection proceeds in an MRX-independent manner at the end predicted to have a 3′ overhang (14, 19, 31). As a control, we expressed Cas9 with the same gRNAs in *mre11Δ* cells and observed the expected persistent two-ended DSB pattern using the same restriction endonuclease digests and hybridization probes (Figure 2C and Figure S2B), confirming that MRX is required to resect both ends of replication-independent DSBs.

**Figure 2.**
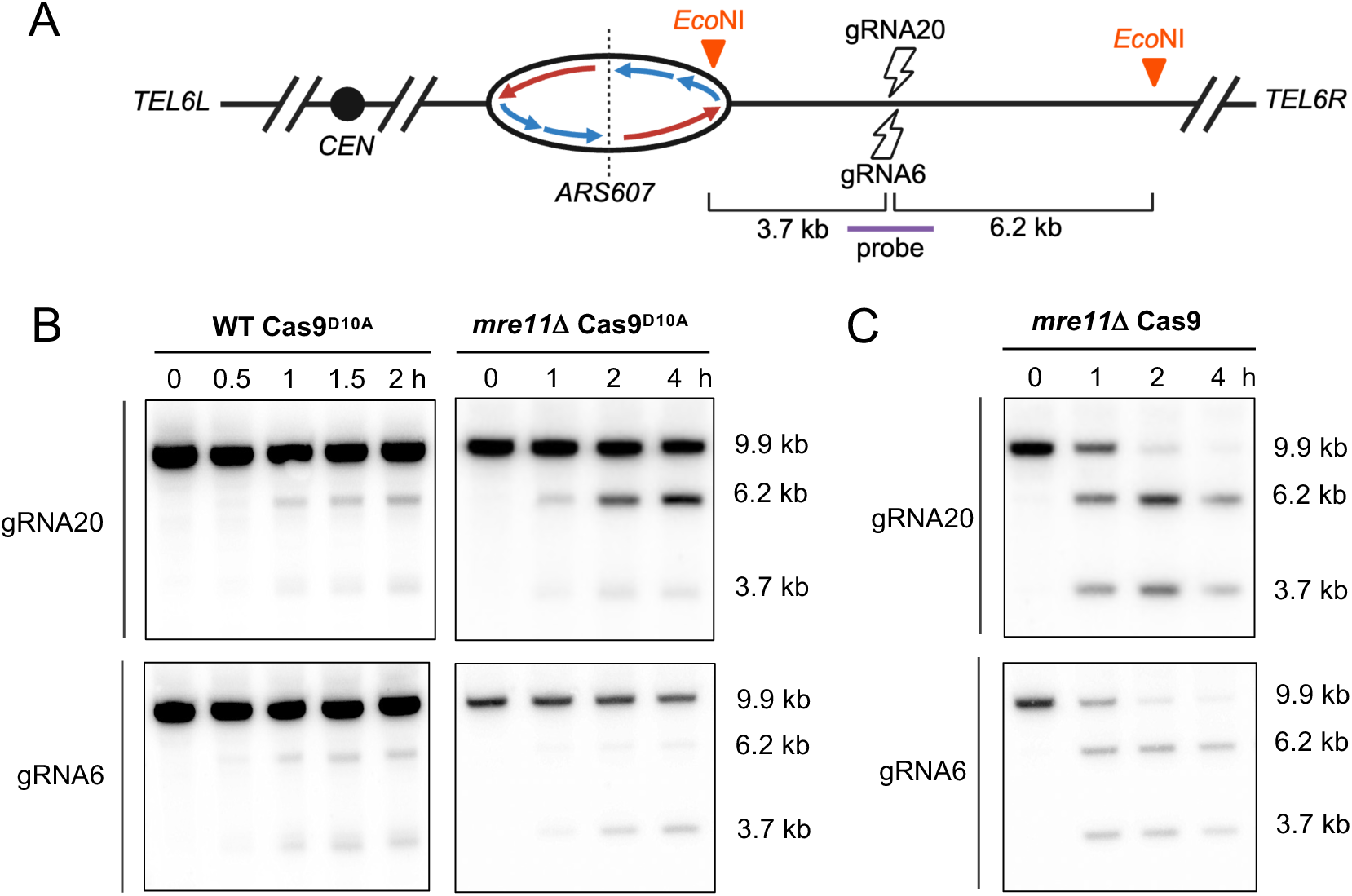
Physical analysis of replication-dependent DSBs in WT and *mre11Δ* cells. **A.** Schematic showing the location of *ARS607*, gRNAs and the sizes of DNA fragments from *Eco*NI digestion of genomic DNA. The 3.7 kb fragment is generated by fork collapse at the nick and 6.2 kb fragment from fork progression beyond the nick or fork convergence from a downstream origin. The 1 kb hybridization probe, indicated by the red line, was designed to hybridize equally with each break fragment. **B.** Southern blots of *Eco*NI-digested DNA before induction and at different time points following expression of Cas9^D10A^ with gRNA 20 or gRNA6 in WT and *mre11Δ* cells. The 9.9 kb fragment corresponds to uncut DNA. **C.** Southern blots of *Eco*NI-digested DNA before induction and at different time points following expression of Cas9 with gRNA 20 or gRNA6 in *mre11Δ* cells.

### MRX activity is required at nearly blunt DSB ends

To directly test this hypothesis, we performed a qPCR-based resection assay measuring the ssDNA fraction in proximity to break sites (Figure S3A) (19, 40). For both gRNA1 and gRNA6, we assessed the ssDNA fraction about 1 kb from each DSB end (labeled CEN and TEL). We strategically measured long-range resection to avoid detecting ssDNA close to the break sites that may result from replication mediated 3′ overhangs. We first verified that Cas9^D10A^ cutting efficiency is comparable and highly efficient 4 hours after nickase induction for all genotypes tested by using a qPCR assay with an amplicon spanning the cut site (Figure S3B, C). To validate that the measured cut fraction was mostly the result of replication-dependent DSBs and not nicks, we arrested cells in G1 and then quantified Cas9^D10A^-induced nicks by their sensitivity to S1 nuclease (Figure S3D, E) (18). We found that prior to S1 nuclease treatment to convert nicks to DSBs (Figure S3D), the cut fraction was negligible 2 hours after nickase induction (Figure S3F). In contrast, treating cells with S1 nuclease resulted in 17% and 24% cleavage products for the gRNA1 and gRNA6 sites, respectively (Figure S3F). This suggests that the cut fraction in cycling cells is mostly comprised of replication-dependent breaks, since the cut fraction reached 59% at the gRNA1 locus and 86% at the gRNA6 locus following two hours after nickase induction in WT cycling cells (Figure S3B, C).

As previously reported (18, 19, 39), *rad51Δ* cells display a hyper resection phenotype at replication-dependent DSBs and were therefore used as controls to detect the maximum ssDNA fraction at the tested loci. In contrast, *exo1Δ sgs1Δ* cells were used as a resection deficient control, as they are expected to have very minimal resection at sites roughly 1 kb distal to a DSB (41). WT cells have intermediate levels of ssDNA because resection intermediates are transient in repair-competent cells (18, 19, 39). We found that blunt ends are processed in a MRX and Exo1/Dna2-Sgs1 dependent manner (Figures 3A, 3C). In contrast, *mre11Δ* cells displayed a much smaller resection defect compared to *exo1Δ sgs1Δ* cells at ends predicted to bear 3′ overhangs (Figures 3A, 3C). While the ssDNA fraction in *mre11Δ* is lower than *rad51Δ* cells at 3′ overhangs, it is apparent that resection can proceed in a mostly MRX independent manner. Since MRX is also known to physically recruit Dna2-Sgs1 and Exo1 to DSBs (36, 42), it is possible that the slight reduction in ssDNA in *mre11Δ* cells at ends with 3′ overhangs compared to *rad51Δ* levels is due to decreased activity of the long-range resection pathways. Taken together, these data further support the model that replication-dependent DSBs are formed asymmetrically because of replication fork dynamics and demonstrate that resection can occur in an MRX independent fashion at 3′ overhang ends of these breaks.

**Figure 3.**
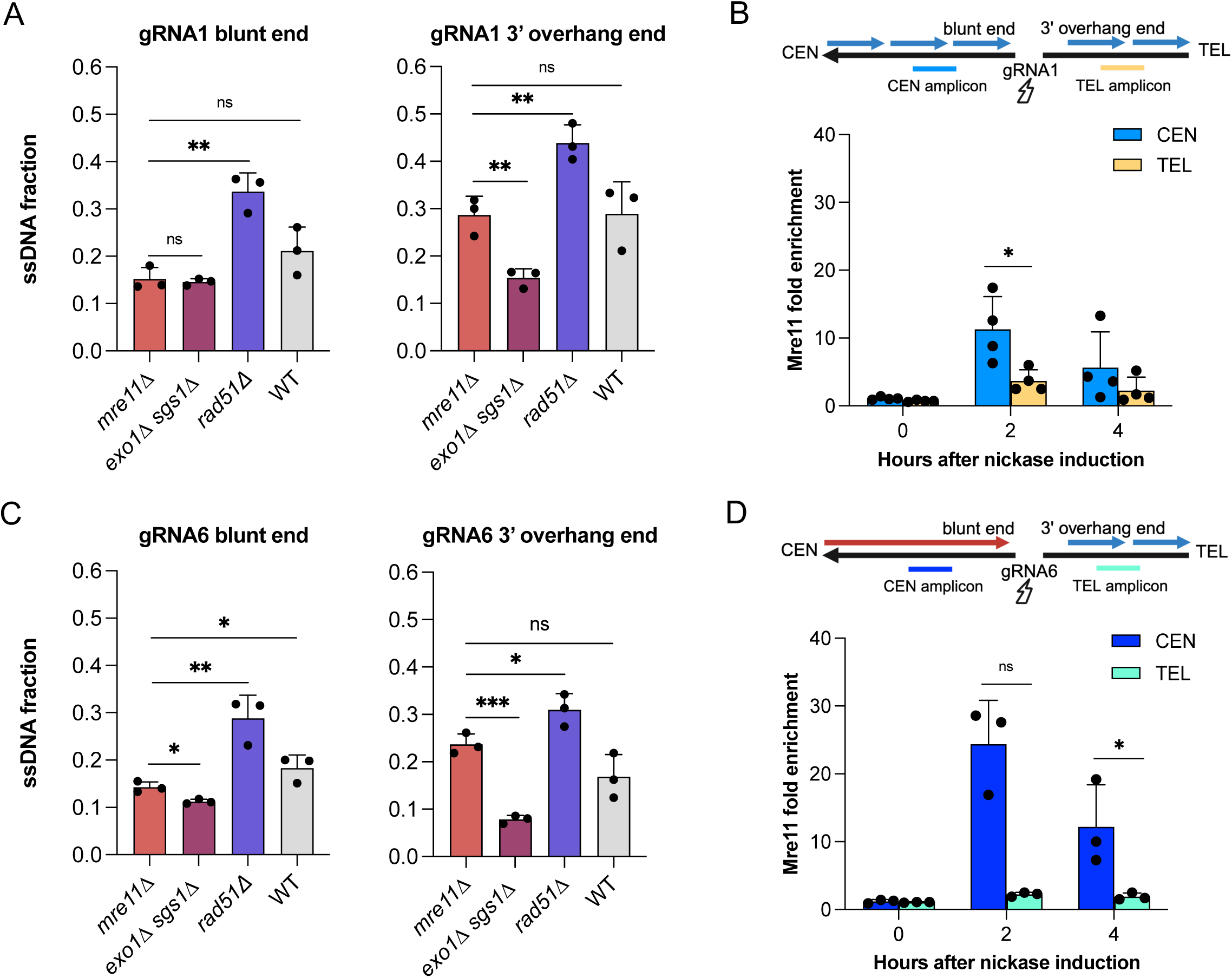
Mre11 preferentially binds to and is required for resection at blunt ends. **A.** Fraction of ssDNA generated at sites approximately 1 kb centromeric and telomeric to the gRNA1 cut site 4 h after Cas9^D10A^ induction assessed with a qPCR-based assay. **B.** Schematic of break orientation at the gRNA1 locus (top), enrichment of Mre11 flanking the gRNA1 locus (bottom) determined by ChIP-qPCR. Amplicons to detect Mre11 binding are located 0.1-0.3 kb centromeric or telomeric to the gRNA1 cut site. **C.** Fraction of ssDNA generated at sites about 1 kb centromeric and telomeric to the gRNA6 cut site 4 h after Cas9^D10A^ induction assessed with a qPCR-based assay. **D.** Schematic of break orientation at the gRNA6 locus (top), enrichment of Mre11 flanking the gRNA6 locus (bottom) determined by ChIP-qPCR. Amplicons to detect Mre11 binding are located 0.2-0.4 kb centromeric or telomeric to the gRNA6 cut site.

To further study the role of MRX in initiating resection, we measured Mre11 binding at replication-dependent DSBs ChIP-qPCR in cycling cells following Cas9^D10A^ induction. We found Mre11 to be more strongly recruited to the ends that are predicted to be blunt than to 3′ overhang ends with gRNA1 and gRNA6 (Figures 3B, 3D). The higher retention of Mre11 at the gRNA6 site could be due to a delay between fork collapse at the leading-strand template nick and arrival of a converging fork to convert the seDSB into a deDSB. In contrast, Mre11 is recruited equally to both sides of a Cas9-induced DSB (Figure S4A). We verified the asymmetrical binding of Mre11 at the gRNA20 locus targeting Cas9^D10A^ to the lagging-strand template (Figure S4B, S4C). We previously reported that a Cas9^D10A^-induced nick is mostly converted to a seDSB using gRNA14 (19), and consistent with this finding we observed enrichment of Mre11 only at the end produced by fork collapse. Nonetheless, the data with the other three gRNAs that generate deDSBs support the notion MRX binding is biased to blunt ends, consistent with end resection being dependent on MRX at these ends.

### Exo1 and Dna2-Sgs1 can process 3′ overhangs in the absence of MRX activity

Finally, we assessed resection dynamics at breaks formed by lagging and leading-strand nicks in resection deficient strains to dissect the relative role of the Exo1/Dna2-Sgs1 resection pathways. At the blunt ends of gRNA1 and gRNA6-generated breaks, resection can proceed efficiently in the *mre11-H125N* strain (Figure 4A, 4B). Upon deletion of *exo1Δ* in the *mre11-H125N* background, we observed a modest reduction in resection (Figure 4A, 4B). This suggests that the Exo1 pathway plays a minor role in the resection of these breaks, and resection is mostly dependent on Dna2-Sgs1 in the absence of MRX nuclease activity. Indeed, the *mre11-H125N sgs1Δ* mutant is almost as resection deficient as the *mre11Δ* strain (Figure 4A, 4B).

**Figure 4.**
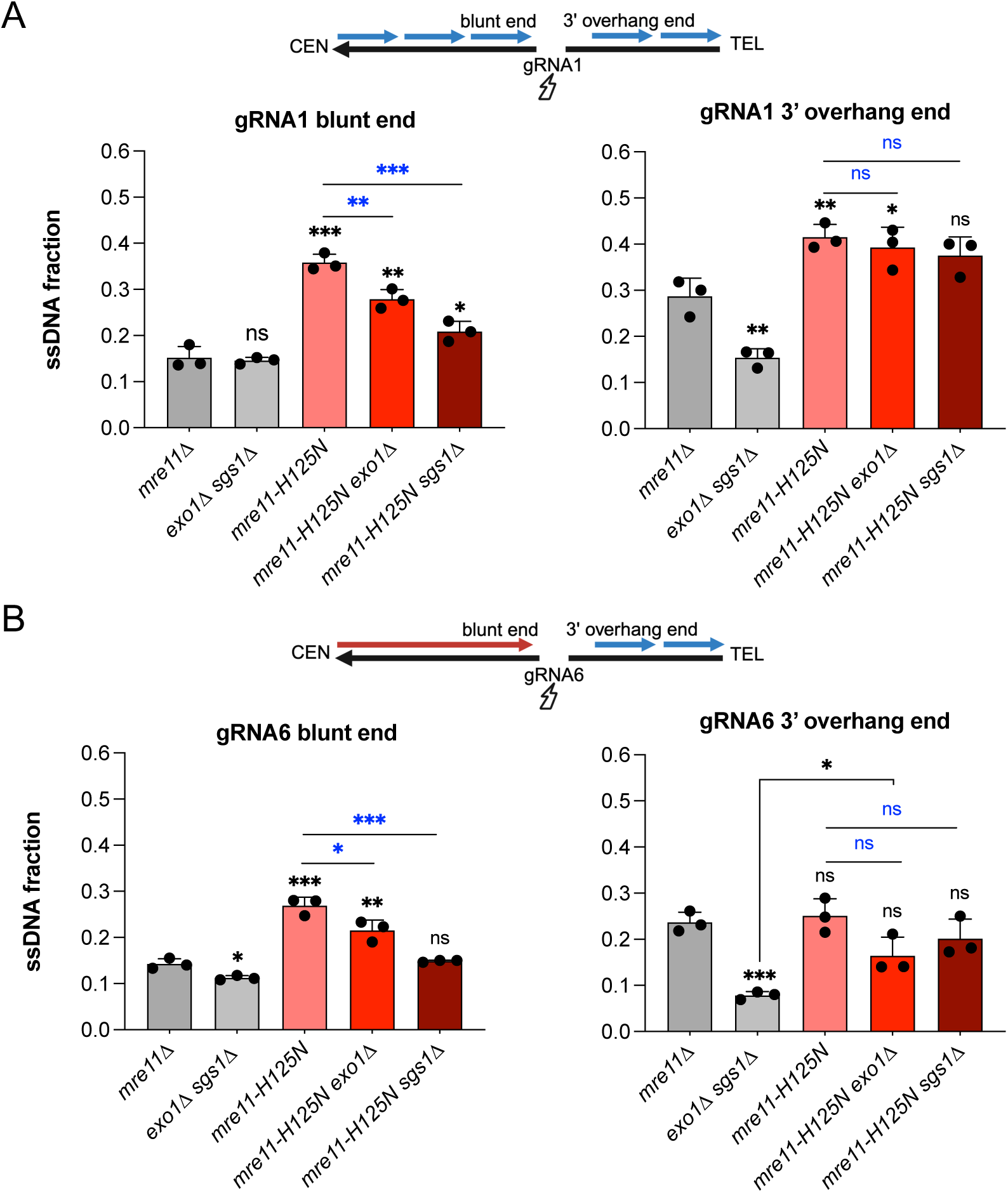
Exo1 and Dna2-Sgs1 can resect DSB ends with 3’ overhangs in the absence of MRX nuclease activity. **A.** Fraction of ssDNA generated at sites approximately 1kb centromeric and 1kb telomeric to the gRNA1 cut site 4 h after Cas9^D10A^ induction assessed by a qPCR-based assay. **B.** Fraction of ssDNA generated at sites approximately 1 kb centromeric and telomeric to the gRNA6 cut site 4 h after Cas9^D10A^ induction assessed by a qPCR-based assay. All significance values are relative to the *mre11Δ* condition using an unpaired T-test. ns p ≥ 0.05, *p ≤ 0.05, **p ≤ 0.01, ***p ≤ 0.001. Significant differences between *mre11-H125N* derivatives are marked with blue asterisks.

In contrast to blunt ends, we observed either a slight increase in resection or an insignificant decrease in resection at 3′ overhang ends in the *mre11-H125N exo1*Δ and *mre11-H125N sgs1*Δ double mutants compared to the *mre11Δ* control (Figure 4A, 4B). The most resection deficient mutant in these experiments was *mre11-H125N exo1Δ* at the gRNA6 locus, however, the ssDNA fraction in this mutant was still more than 2-fold higher than in the *exo1Δ sgs1Δ* mutant (16.4% vs. 7.8%). The resection profiles from the 3’ overhang ends at both guide sites indicate that in the absence of MRX nuclease activity, either the Exo1 or Dna2-Sgs1 pathway can initiate resection. In contrast, Dna2-Sgs1 is required for the processing of blunt ends in the absence of MRX nuclease activity with only a minor contribution by Exo1.

### MRX activity is required for removal of Ku at blunt ends of replication-dependent DSBs

Previous studies have shown that removal of Ku partially suppresses the DNA damage sensitivity and end resection defect of Mre11-deficient cells in an Exo1-dependent manner (36, 37, 43, 44). Furthermore, over-expression of Exo1 suppresses the DNA damage sensitivity of the *mre11Δ* mutant (45–47). To increase Exo1 expression, we used a previously constructed high-copy number plasmid with *EXO1* expressed from its native promoter (37) We found that over-expressing Exo1 rescued the sensitivity of *mre11Δ* cells to Cas9^D10A^-induced nicking with gRNA1 by ∼100-fold compared to the empty vector control (Figure 5A). Surprisingly, Exo1 over-expression was less effective in relieving the sensitivity *mre11Δ* cells to induction of Cas9^D10A^ with gRNA6 (Figure 5A). This finding suggests that leading strand collapses are processed in a distinct way from lagging strand DSBs. To confirm this finding, we also measured the viability of *mre11Δ* and *mre11Δ* cells +/-Exo1 overexpression in response to gRNA14 or gRNA20 expression with Cas9^D10A^. These guides convey less of a growth defect to *mre11Δ* cells than gRNAs 1 and 6, however, we still found that the lagging strand DSB showed less of a dependence on MRX for repair under Exo1 overexpression conditions when compared to the leading strand collapse (Figure S6A). Similarly to Exo1 overexpression, loss of Ku showed a stronger suppression of *mre11Δ* sensitivity to gRNA1 than to gRNA6 (Figure 5B).

**Figure 5.**
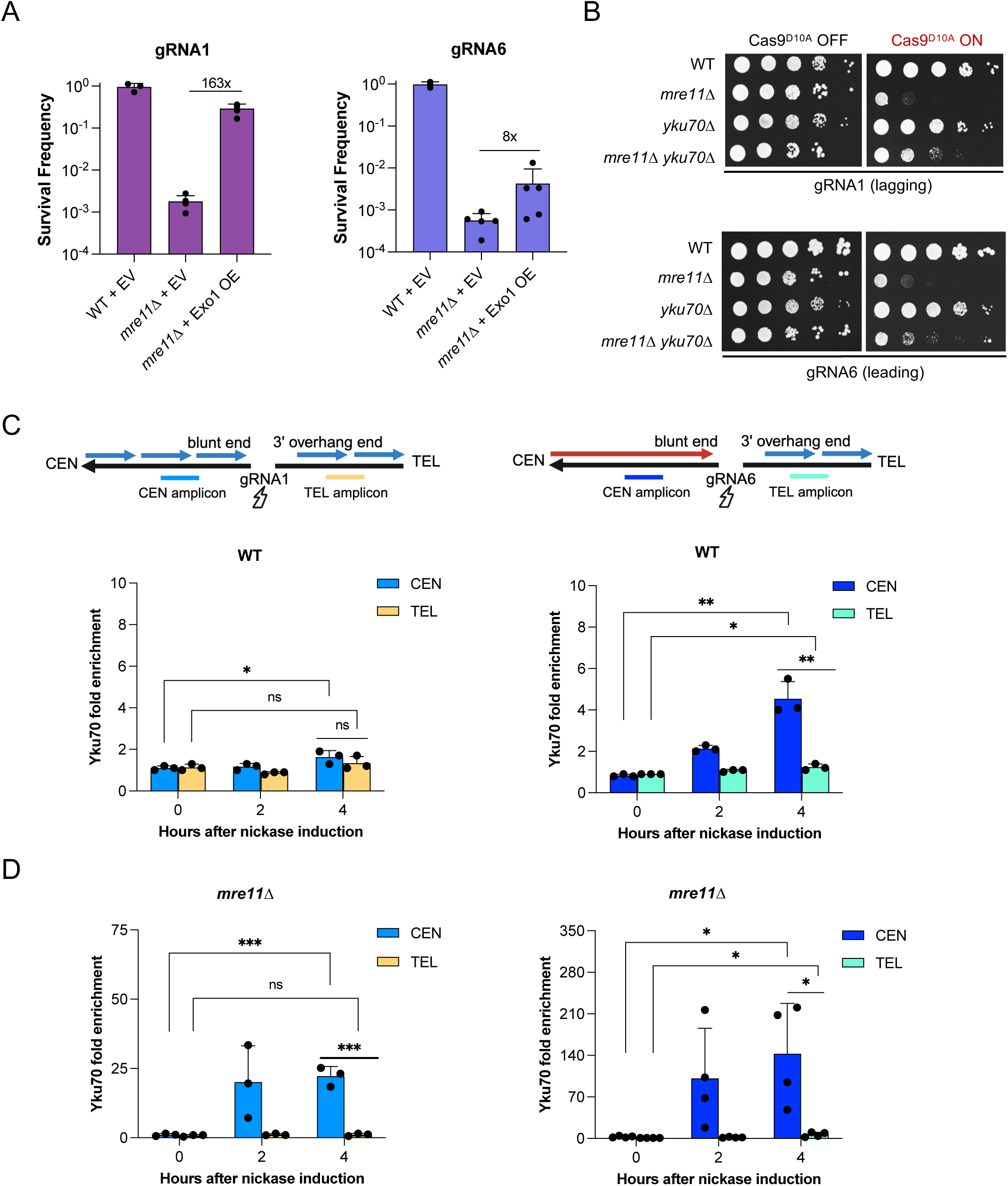
Lagging strand breaks are more accessible to Exo1. **A.** Fraction of *mre11Δ* and *mre11Δ* + Exo1 OE cells that survived expression of Cas9^D10A^ with gRNA1 (left) or gRNA6 (right). **B.** Serial dilutions of the indicated strains expressing gRNA1 or gRNA6 were spotted on medium -/+ β-estradiol to induce Cas9^D10A^ expression. **C.** Schematic of break orientation at the gRNA1 or gRNA6 locus (top), enrichment of Yku70 at sequences flanking the gRNA1 or gRNA6 site in WT cells at 0, 2 or 4 h post Cas9^D10A^ induction determined by ChIP-qPCR. **D.** Enrichment of Yku70 at sequences flanking the gRNA1 or gRNA6 locus in *mre11Δ* cells. Amplicons to detect Yku70 binding are located 0.1-0.3 kb centromeric or telomeric to the gRNA1 cut site, or 0.2-0.4 kb centromeric or telomeric to gRNA6. All significance values calculated using an unpaired t-test. ns p ≥ 0.05, *p ≤ 0.05, **p ≤ 0.01, ***p ≤ 0.001.

Since Exo1 functions redundantly with Dna2-Sgs1 in long-range resection, we also tested whether overexpressing Dna2 could rescue the growth defect of *mre11Δ* cells in response to expression with gRNA1 or gRNA6. In contrast to Exo1 overexpression, we found that Dna2 expression compounded the repair defect of *mre11Δ* cells with both gRNAs (Figure S6B). This finding agrees with previous work demonstrating that Dna2 overexpression sensitizes cells to DNA damaging agents such as camptothecin (48). Similarly, overexpressing a nuclease dead allele of Exo1, *exo1^D173A^*, also exacerbated the repair defect of *mre11Δ* cells in the gRNA2 condition (Figure S6B), likely due to the nuclease-dead protein blocking DSB ends from potential resection by endogenous Exo1 or Dna2-Sgs1.

To test whether Ku prevents access to Exo1 at the leading strand collapse, we performed ChIP-qPCR for Yku70 on WT and *mre11Δ* cells targeted with both gRNA1 and gRNA6. We found that Ku plays a relatively minimal role at the gRNA1 locus in WT cells with a peak enrichment of 1.9-fold, while Ku is strongly enriched at the blunt end of the gRNA6 locus leading strand collapse with a peak enrichment of 4.5-fold (Figure 5C). Notably, we observed minimal Ku enrichment at the 3′ overhang end flanking the gRNA1 and gRNA6 breaks (Figure 5C). In the absence of Mre11, Ku enrichment at blunt ends increased by 10-fold at both leading and lagging strand breaks, but there was still little Ku binding at 3′ overhang ends (Figure 5D). We also measured Ku enrichment at a Cas9 DSB targeted with gRNA1. As expected, and in contrast to Cas9^D10A^ induced breaks, we found strong, symmetrical Yku70 enrichment to both DSB ends (Figure S6C). The asymmetrical binding of Ku to replication-dependent DSBs likely contributes to the failure to repair these breaks by NHEJ (19).

## Discussion

Here we investigated resection dynamics at DSBs formed by replication fork collision with a nick on the leading or lagging-strand template. In brief, we find that the MRX complex is only required to initiate resection at deDSB ends that are predicted to be blunt, or nearly blunt, but not at ends predicted to have 3′ overhangs resulting from lagging-strand synthesis. Consistently, we observe higher Mre11 binding to blunt ends of deDSBs than to 3′ overhang ends. We suggest that Exo1 and Dna2-Sgs1 can directly resect ends with long (∼70 nt) 3′ overhangs, bypassing the need for resection initiation by MRX. Although it has been shown that resection of ends produced by ionizing radiation or endonucleases is uncoordinated in the absence of the MRX complex (49, 50), this is likely due to the low efficiency of resection initiation at each end of a DSB, rather than to different end structures. Our findings have some parallels with telomeres, where it has been shown that resection can proceed in the absence of MRX at lagging-strand telomeres, but not at leading-strand telomeres (51). The asymmetrical recognition and processing of the ends of replication-dependent deDSBs agrees with *in vitro* studies showing that replication fork collapse at nicks on leading and lagging strand templates results in seDSBs with different end structures (31).

The finding that resection can proceed at one end of replication-dependent deDSBs in the absence of MRX raises the question of why the *mre11*Δ mutant shows such high sensitivity to nickase expression. It is possible that the 3′ overhang end can invade the sister chromatid to initiate repair synthesis but in the absence of timely second-end capture, the cells cannot complete repair. The other possible role for MRX is to maintain sister-chromatid cohesion during repair of replication-dependent DSBs. Previous studies have shown an increase in spontaneous and nick-induced inter-chromosomal recombination in the absence of the MRX complex in yeast, likely because of a sister-chromatid cohesion defect (19, 52). Nick-induced unequal sister chromatid recombination between closely linked repeats is also less impacted by loss of Mre11 (27, 38). How MRX achieves sister chromatid tethering is currently unknown. Several groups argue that the Rad50 subunit of MRX might tether sister chromatids at the site of a DSB through its long coiled-coil domains, analogous to cohesin, (52–56). Others propose that through initiating end resection, MRX assists in *de novo* cohesin loading which indirectly stimulates sister chromatid tethering (57, 58). Resolving whether MRX plays a direct physical role maintaining sister chromatid tethering, or, if MRX initiated resection stimulates sister chromatid tethering would help better define how replication-dependent DSBs are repaired.

Additionally, we find that Ku binds blunt ends of replication-dependent DSBs, and that MRX is required to remove Ku from these ends. This finding is consistent with studies showing delayed clearance of Ku from endonuclease-induced DSBs in the absence of MRX (59). An alternative explanation for this finding is that MRX physically competes with Ku for access to DNA ends. We did not detect Ku enrichment at 3′ overhang ends, even in the absence of MRX. Our results contrast with telomeres where Ku has been found to associate with both leading and lagging-strand telomeres (51). If the 3′ overhang ends are rapidly resected by Exo1 and/or Dna2-Sgs1 then the long tracts of ssDNA would be poor substrates for Ku binding. Furthermore, RPA bound to ends with ssDNA overhangs has been shown to prevent access to Ku (60). Since the 3′ overhang ends of replication-dependent DSBs would be bound by RPA, they would not be available for Ku to bind even prior to long-range resection.

Unexpectedly, we find higher enrichment of Ku at the site of a leading strand collapse, as opposed to a nearly blunt end generated by termination of an Okazaki fragment at the site of a nick. The reason for this difference is not immediately obvious. It is possible that during Okazaki fragment termination, DNA polymerase 8 (Pol8) remains bound to the site of the DSB. In this scenario, Ku would have to compete with Pol8 for DSB binding. In contrast, leading strand synthesis is carried out by Polε, which is coupled to the CMG helicase (61). Since *in vitro* work demonstrates that the CMG helicase dissociates from DNA when a nick is induced on a leading-strand template (18, 31), we propose that these ends might be less protected than nearly blunt ends resulting from lagging strand synthesis. This could result in higher Ku enrichment at a leading strand collapse than to blunt ends of lagging strand DSBs. In support of this idea, Exo1 overexpression more robustly rescues the growth defect of *mre11Δ* cells challenged with a lagging strand DSB compared to a leading strand collapse, suggesting that Ku plays less of an inhibitory role to resection at lagging strand DSBs. Another possibility is that the blunt ends of breaks formed by nicks on the lagging-strand template have a different structure to the end formed by CMG run off at a nick on the leading-strand template. The *in vitro* study mapping ends generated at pre-formed nicks on plasmid templates detected only seDSBs (31); therefore, the structure of the end produced by polymerase run off at a nick on the lagging strand template to form a deDSBs is uncharacterized. Since Exo1 preferentially degrades 5′ recessed ends, has reduced activity at blunt ends, and low activity at ends with 5′ overhangs (62), the end produced by a leading strand collapse would be a poor substrate for Exo1.

The physiological relevance of Ku binding to one end of replication-dependent DSBs is unclear as these ends are not substrates for NHEJ repair (19, 20, 23, 27). It is possible that Ku plays a role in synchronizing resection at replication-dependent DSBs to minimize potentially mutagenic ssDNA tracts. Consistent with this idea, work in *S. pombe* has demonstrated that the mutagenic break-induced replication (BIR) pathway is more active in the absence of the Ku complex (20). Finally, since the Ku protein block has been shown to stimulate MRX endonuclease activity *in vitro* (8, 63), Ku may function at asymmetric DSB ends to ensure that resection proceeds in a mostly symmetric manner. One potential hazard of Ku binding to one end of replication-dependent DSBs is the possibility of joining the blunt ends of two independent replication dependent DSBs, resulting in chromosome rearrangements.

In summary, this work details how resection proceeds at DSBs generated during DNA replication. While our findings are in line with some aspects of resection regulation at canonical DSBs, we also find key differences in repair factor binding and nuclease usage at each end of replication-dependent deDSBs. It is interesting to note that although BRCA1 is required for resection of endonuclease-induced DSBs in mammalian cells, it is dispensable for resection of replication-dependent DSBs (18, 27). Since Cas9 nickases, such as Cas9^D10A^, are currently being studied for their gene editing potential (64, 65), the finding that nicks can trigger BIR (22, 27, 28) raise the possibility of unintended off target effects. In future work, it would be interesting to investigate mutagenic outcomes of Cas9 nickase-induced breaks, and the genetic backgrounds that might be prone to these negative outcomes.

## Materials and Methods

### Media and yeast strains

Complete yeast media contained 1% yeast extract, 2% peptone, 10 µg/mL adenine, and 2% glucose (YPAD) as a carbon source. Synthetic media contained 1X yeast nitrogen base, 1X amino acid dropout mix, and 2% glucose. For β-estradiol induction of Cas9^D10A^, a 10 mM stock of β-estradiol was diluted and spread on plates or added to liquid media for a final concentration of 2 µM.

All yeast strains are in the *RAD5*-corrected W303 background (*leu2-3*,*112 trp1-1 can1-100 ura3-1 ade2-1 his3-11,15*) unless otherwise noted and are listed in Supplementary Table 1. Strains were constructed by standard genetic methods. Lithium acetate transformation was used to introduce plasmids or deletion cassettes containing a marker of choice and homology arms flanking the gene to be deleted. Other strains were made by genetic cross, followed by tetrad dissection and marker selection.

### Plasmids

#### Cas9 and gRNA plasmids

Plasmids expressing FLAG tagged Cas9 (pLS504) or Cas9^D10A^ (pLS503) were generated in the pRG203MX and integrated into the genome as previously described (19). For strains used for Yku70-FLAG ChIP, we derived plasmids without a FLAG tag fused to Cas9^D10A^ or Cas9 (pLS755 and pLS758 respectively) by amplifying the entire pLS503 or pLS504 plasmid except the 3xFLAG tag by PCR (see Table S2 for primers) and ligating the resulting linear product with T4 DNA ligase after T4 polynucleotide kinase treatment. Guide RNAs were expressed in a pRG205MX (*LEU2*) integrating plasmid backbone (pLS505) as previously described (see Table S3) (19).

Episomal centromeric plasmids were previously constructed that express both Cas9^D10A^ and either gRNA1, gRNA6, gRNA14, or gRNA20 (pLS627, pLS633, pLS733, and pLS735 respectively) (19). Full plasmid sequences are available upon request.

#### Overexpression plasmids

Overexpression plasmids were derived from the high copy number vector pRS426 (*URA3*). Full plasmid sequences are available upon request.

### Spot assays for assessing growth sensitivity of Cas9^D10A^ strains

For all spot assays except those testing Exo1 over-expression, Cas9^D10A^ and the relevant gRNA were co-expressed from a centromeric plasmid. Single colonies were used to inoculate 2 mL SC-Leu media and cultures were grown overnight at 30°C while shaking. The following day, 10-fold serial dilutions were spotted on media with or without 2 µM β-estradiol. Plates were grown for three days at 30°C, unless otherwise noted, after which plates were imaged with a desktop scanner.

### Survival assays for overexpression experiments

All survival assays with Cas9^D10A^ were done in strains in which the relevant gRNA was integrated into the genome and grown on SC-URA media to maintain episomal overexpression vectors. Fresh single colonies from each strain were picked and grown for four hours in SC-URA liquid media. Cells were then diluted and plated in technical replicate on SC-URA ± β-estradiol. Colonies were allowed to grow for five days at 30°C, then counted. The survival frequency was calculated by the formula:

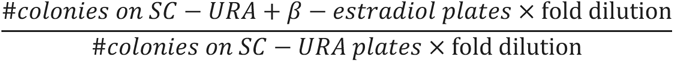

### Physical analysis of Cas9^D10A^-induced breaks by Southern blotting

#### Induction of Cas9^D10A^ in asynchronous cultures

Strains with integrated Cas9^D10A^ and gRNA cassettes were used for physical analysis of replication-dependent DSBs. For Cas9^D10A^ induction, single colonies of relevant strains were used to inoculate 5 mL YPAD cultures that were grown overnight at 30°C. The following day, cultures were diluted to an OD_600_ of 0.4 in YPAD and grown for 1 hour. 1 x 10^8^ cells were collected before induction (t_0_) and then β-estradiol was added to the remaining culture at a final concentration of 2 µM to induce Cas9^D10A^. The same number of cells were collected at each timepoint, sodium azide was added to 0.1% final, cells were collected by centrifugation at 3,000 rpm for 5 min at 4°C, cell pellets were then washed with dH_2_O, centrifuged at 2,000 x g for 2 min at 4°C in a microcentrifuge and cell pellets were stored on ice prior to embedding in agarose plugs. Cells were collected at 0-, 1-, 2-, and 4-hour points and agarose plugs were prepared and processed as previously described (19, 66), followed by Southern blotting (19).

### Flow cytometry

Two volumes of 95% ethanol were added to cultures containing 1 x 10^7^ cells, vortexed, and stored at -20°C freezer overnight. The following day, cells were collected by centrifugation at 2,000 x g for 3 min at room temperature, resuspended twice in 500 µL of 50 mM Tris-HCl pH7.5, incubated for 10 min at room temperature and collected at 2,000 x g for 3 min. 500 µL of RNAse A solution (1 mg/mL RNAse A in 50 mM Tris-HCl pH7.5, 7.5 mM NaCl) was added to each pellet, and incubated at 37°C overnight. The following day, cells were collected, resuspended in 500 µL of 1 mg/mL Proteinase K in 50 mM Tris-HCl pH7.5, and incubated at 50°C for 1 h. Cells were collected, resuspended in 500 µL of 200 mM Tris-HCl pH7.5, 200 mM NaCl, 80 mM MgCl_2_, stained with 300 nM SYTOX Green, and sonicated briefly. Fluorescence was measured using an LSR-II (Becton Dickinson) and FACSDiva (Becton Dickinson) software using a 488 nm blue laser and 525/50 nm emission filter. Data analysis was performed using FlowJo_v10 (Becton Dickinson Biosciences) software.

### Quantitative PCR (qPCR)-based resection assay

Resection was measured using a qPCR-based assay as previously described (67) with the following modifications. ∼10^8^ cycling cells were collected at the indicated time intervals after β-estradiol was added to a final concentration of 2 µM for Cas9^D10A^ induction. Per sample and amplicon, reactions were performed in technical triplicates containing 4.4 µL digested or undigested genomic DNA, 300 nM of each forward and reverse primer (listed in Table S4), and 1x SsoAdvanced Universal SYBR Green Supermix in a total volume of 10 μL and analyzed them on a CFX384 Real-Time System with 10 minutes initial denaturation at 95°C, followed by 40 cycles of 1 minute denaturation at 95°C and 1 minute annealing and extension at 58°C. The fraction of ssDNA, *f_ssDNA_*, at the restriction enzyme recognition site was calculated as previously described (19) with the following formula:

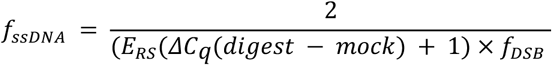

where *E_RS_* is the primer efficiencies for the RS amplicon, *ΔC_q_(digest-mock)* is the difference between quantification cycles for the digested minus the mock-digested sample, and *f_DSB_* is the fraction of genomes where Cas9^D10A^ cleaved the target locus. *f_DSB_* for a locus cleaved by Cas9^D10A^ can be calculated as:

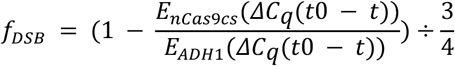

where *E_nCas9cs_* and *E_ADH1_* are the primer efficiencies for the Cas9^D10A^-cut site and ADH1 amplicons and *ΔC_q_(t_0_ - t)* is the difference between the quantification cycles at the first time point (before β-estradiol induction of Cas9^D10A^) and the evaluated time point.

### ChIP-qPCR to measure protein recruitment to replication-dependent DSBs

Protein recruitment to replication-dependent DSBs was assessed with a ChIP-qPCR assay. 25 mL cycling cells at a density of OD_600_ = 0.4 were collected at the indicated time intervals after β-estradiol was added to a final concentration of 2 µM for Cas9^D10A^ induction. 37% formaldehyde was added to a final concentration of 1% and incubated at RT for 15 minutes with shaking. The crosslinking reaction was quenched with 2.5 M glycine, followed by washing with HEPES buffered saline and ChIP lysis buffer (50 mM HEPES, 140 mM NaCl, 1 mM EDTA, 1% IGEPAL CA-630, 0.1% Sodium deoxycholate).

Cells were lysed in ChIP lysis buffer containing 1 mM PMSF using a FastPrep-24 (MP Biomedicals) protocol, run for 30 seconds at 6.5 m/s 4 times. Lysate was placed on ice between each cycle for 45 seconds. Chromatin was fragmented with a QSonica Q800R sonicator for 10 cycles with an amplitude of 65% for 30 seconds. After clearing the lysate, IP samples were incubated with the appropriate antibody pre-incubated with A/B Dynabeads in lysis buffer at a concentration of 1:20 for 3 hours. Control input DNA samples were kept on ice during immunoprecipitation of the remaining DNA. Immunoprecipitated DNA was washed and reverse crosslinked at 65°C for 16 hours.

Input and IP DNA was purified using the Monarch PCR and DNA cleanup kit (NEB). qPCR reactions were performed in the same manner as described in the resection assay with the following modification to the CFX384 Real-Time System protocol. qPCR reactions were run with a 3 minute initial denaturation at 95°C, followed by 40 cycles of 15 second denaturation at 95°C and 30 second annealing and extension at 60°C. Fold enrichment is defined as 2^ΔΔCq^, where ΔΔCq is defined as:

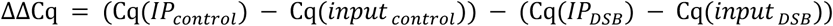

### Quantification and statistical analysis

Quantification and statistical analysis were done using PRISM (GraphPad). ChIP-qPCR experiments were performed at least two times and statistical significance was determined by a two-tailed Student’s t test. For ChIP-qPCR and survival assays, error bars indicate standard deviation. The two-tailed Student’s t test was also used for all survival assays, with the following symbols to indicate significance thresholds: ns = not significant p > 0.05, *p < 0.05, ***p < 0.001, ****p < 0.0001.

## Acknowledgements

We thank all the members of the Symington lab for review of the manuscript. This work was supported by grants from the National Institutes of Health (R35 GM126997 to L.S.S. and T32 CA265828 to M.J.J.). Schematics shown in the figures were generated in BioRender.

---

**Figure S1.**
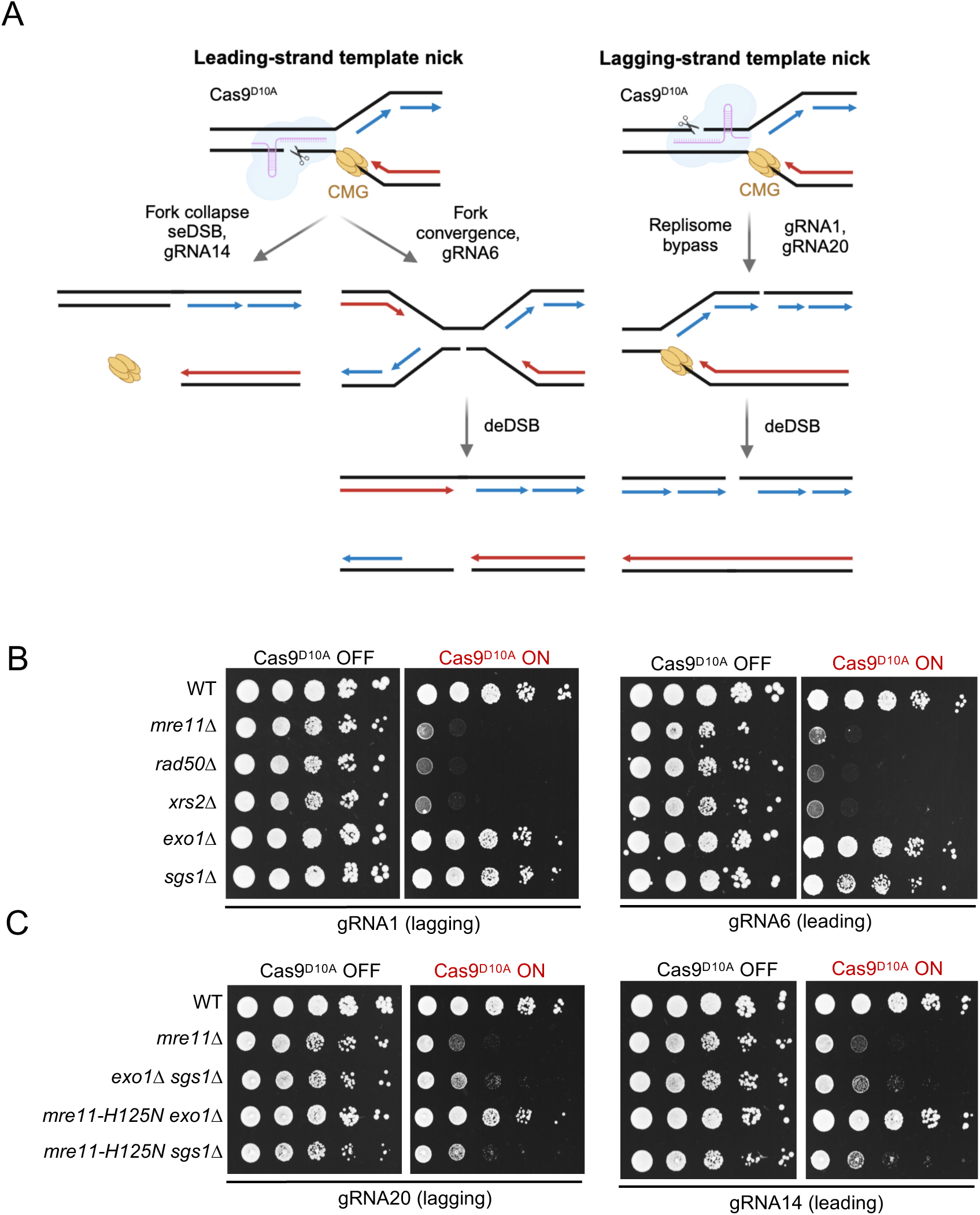
Cas9^D10A^ induced replication-dependent DSBs require end resection for repair. **A.** Model for generation of replication-dependent DSBs at nicks induced by Cas9^D10A^ on the leading or lagging strand template. **B.** Ten-fold serial dilutions of of the indicated strains expressing gRNA1 or gRNA6 spotted on medium -/+ β-estradiol to induce Cas9^D10A^ expression. **C.** Ten-fold serial dilutions of of the indicated strains expressing gRNA20 or gRNA14 spotted on medium -/+ β-estradiol to induce Cas9^D10A^ expression.

**Figure S2.**
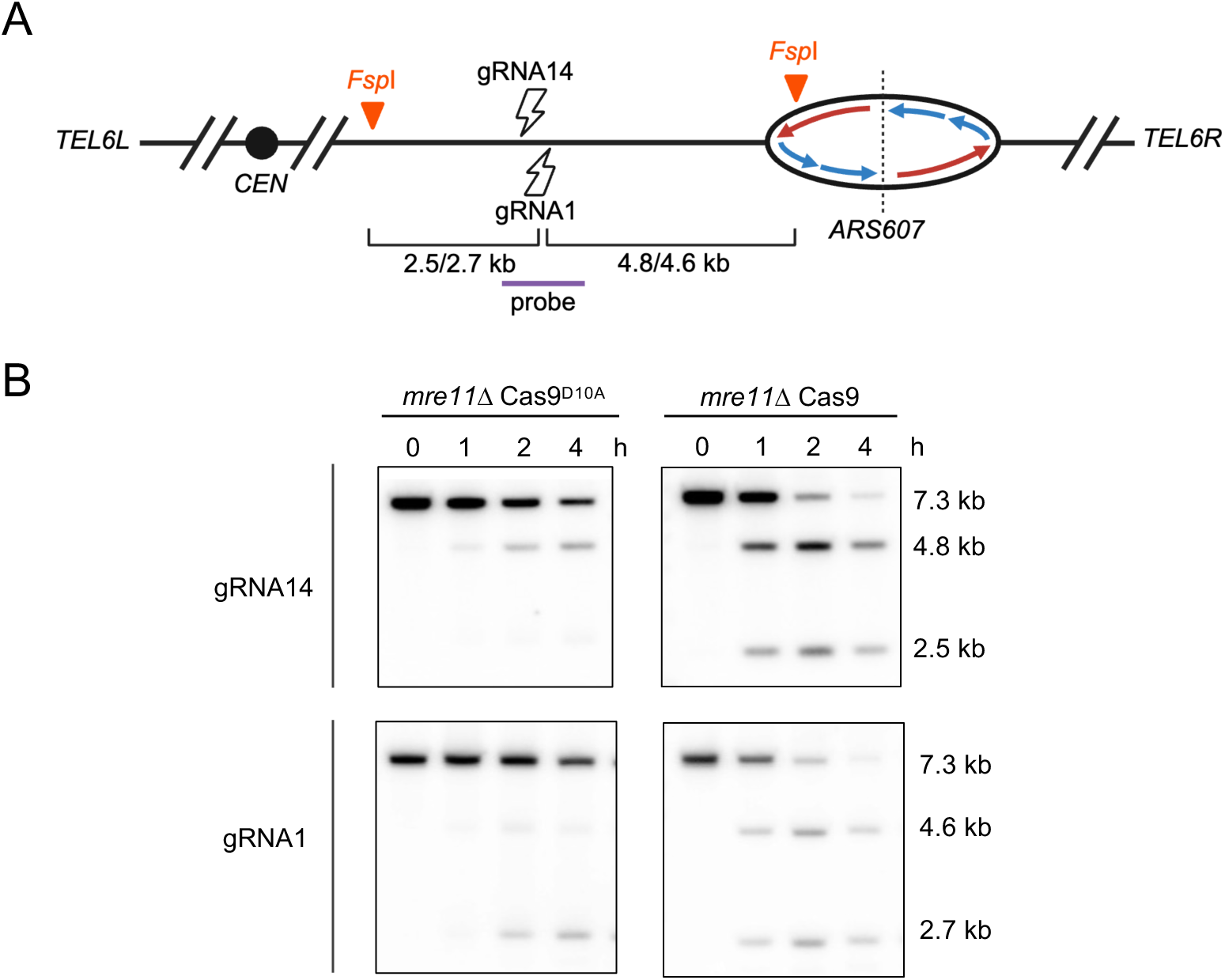
Physical analysis of replication-dependent DSBs in *mre11Δ* cells. **A.** Schematic showing the location of *ARS607*, gRNAs and the sizes of DNA fragments from *Fsp*I digestion of genomic DNA. gRNA14 produces fragments of 2.5 and 4.8 kb with Cas9, and gRNA1 produces fragments of 2.7 and 4.6 kb. Fork collapse at the nick would produce fragments of 4.8 or 4.6 kb with gRNA14 or gRNA1, respectively. A deDSB is indicated by the 2.5 or 2.7 kb fragment. **B.** Southern blots of *Fsp*I-digested DNA before induction and at different time points following expression of Cas9^D10A^ or Cas9 with gRNA14 or gRNA1 in *mre11Δ* cells. The 7.3 kb fragment corresponds to uncut DNA.

**Figure S3.**
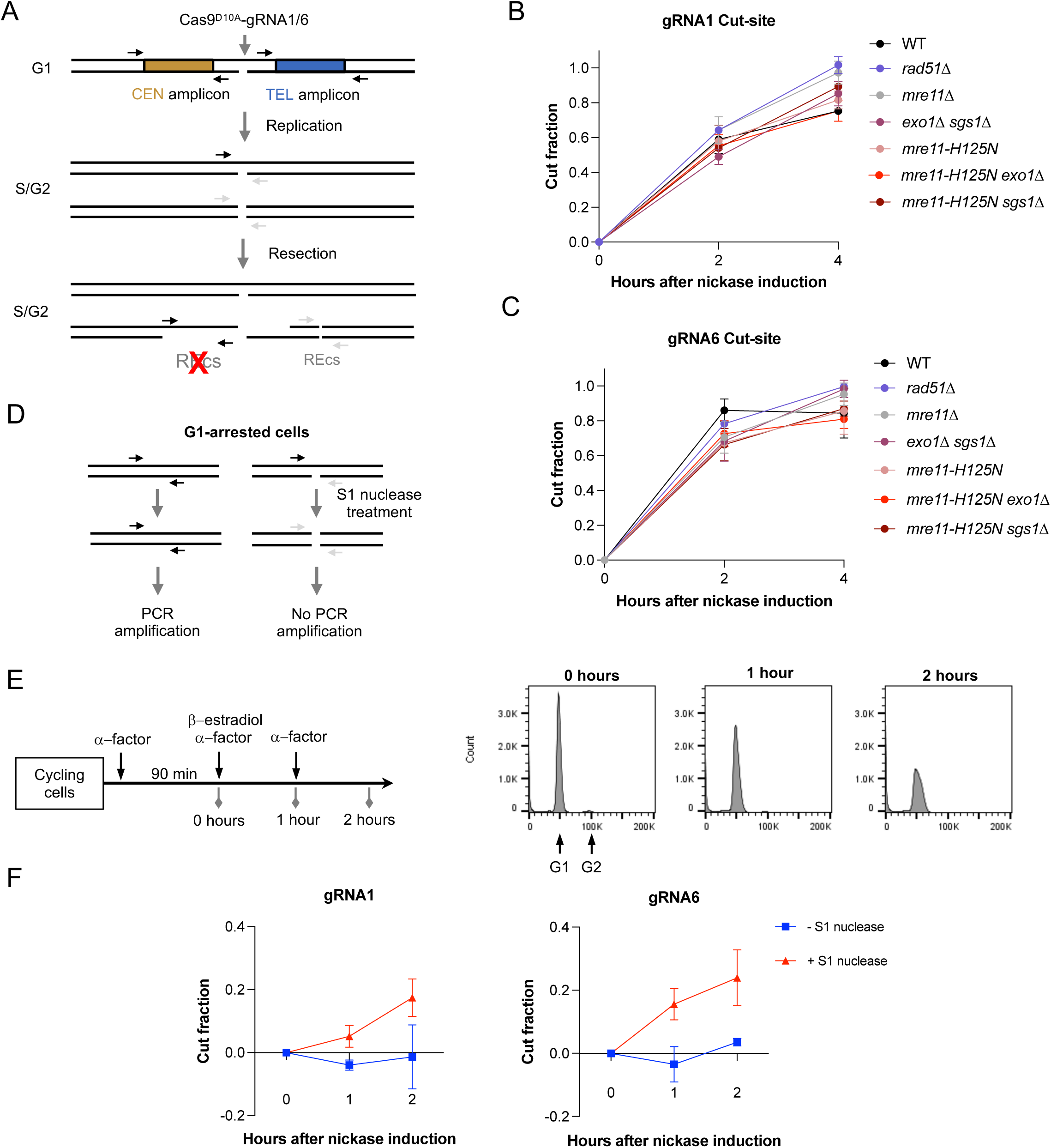
Cut fractions at the gRNA1 and gRNA6 sites after nickase induction. **A.** Schematic showing the location of the gRNA1 or gRNA6-induced nicks and primers flanking restriction endonuclease sites for the qPCR resection assay. After replication, the nicked strand is converted to a deDSB that undergoes end resection to generate ssDNA. The cut fraction is calculated with primers flanking the cut-site, taking into account that nicking can continue in the replicated template. Unresected genomic DNA can be cleaved by restriction endonucleases preventing PCR amplification, whereas ssDNA is resistant to digestion and can be amplified. Cutting efficiency at the gRNA1 (**B**) and gRNA6 (**C**) sites at 2 and 4 hours post induction**. D.** Schematic demonstrating how S1 nuclease digest can convert nicks to DSBs preventing PCR amplification using primers flanking the cut site. Prior to S1 treatment, there would be one intact and one nicked strand and thus result in only a 50% decrease in the PCR product. **E.** Schematic showing G1 arrest and nickase induction growth regime (left). FACS plots of synchronized *bar1Δ* cells collected at 0, 1, and 2 hours after nickase induction. **F.** Cutting efficiency at the gRNA1 and gRNA6 sites at 1 and 2 hours post nickase induction in *bar1Δ* cells.

**Figure S4.**
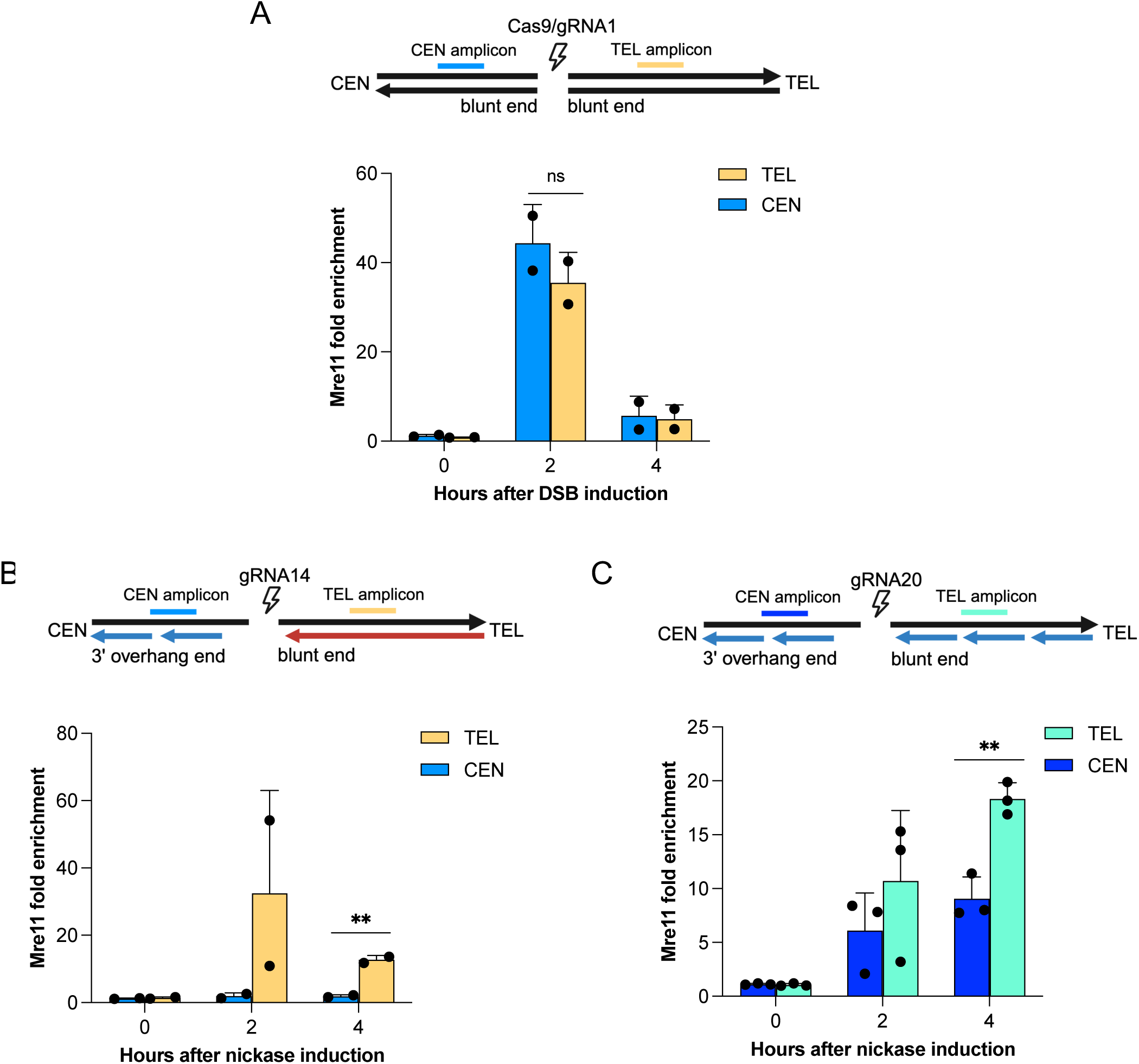
Mre11 enrichment in other break contexts. **A.** Schematic of break orientation at the gRNA1 locus targeted by Cas9 (top), enrichment of Mre11 flanking the gRNA1 locus targeted by Cas9 (bottom). Amplicons to detect Mre11 binding are located 0.1-0.3 kb centromeric or telomeric to the gRNA1 cut site. **B.** Schematic of break orientation at the gRNA14 locus (top), enrichment of Mre11 flanking the gRNA14 locus (bottom). Amplicons to detect Mre11 binding are located 0.1-0.3 kb centromeric or telomeric to the gRNA1 cut site. **C.** Schematic of break orientation at the gRNA20 locus (top), enrichment of Mre11 flanking the gRNA20 locus (bottom). Amplicons to detect Mre11 binding are located 0.1-0.3 kb centromeric or telomeric to the gRNA1 cut site.

**Figure S5.**
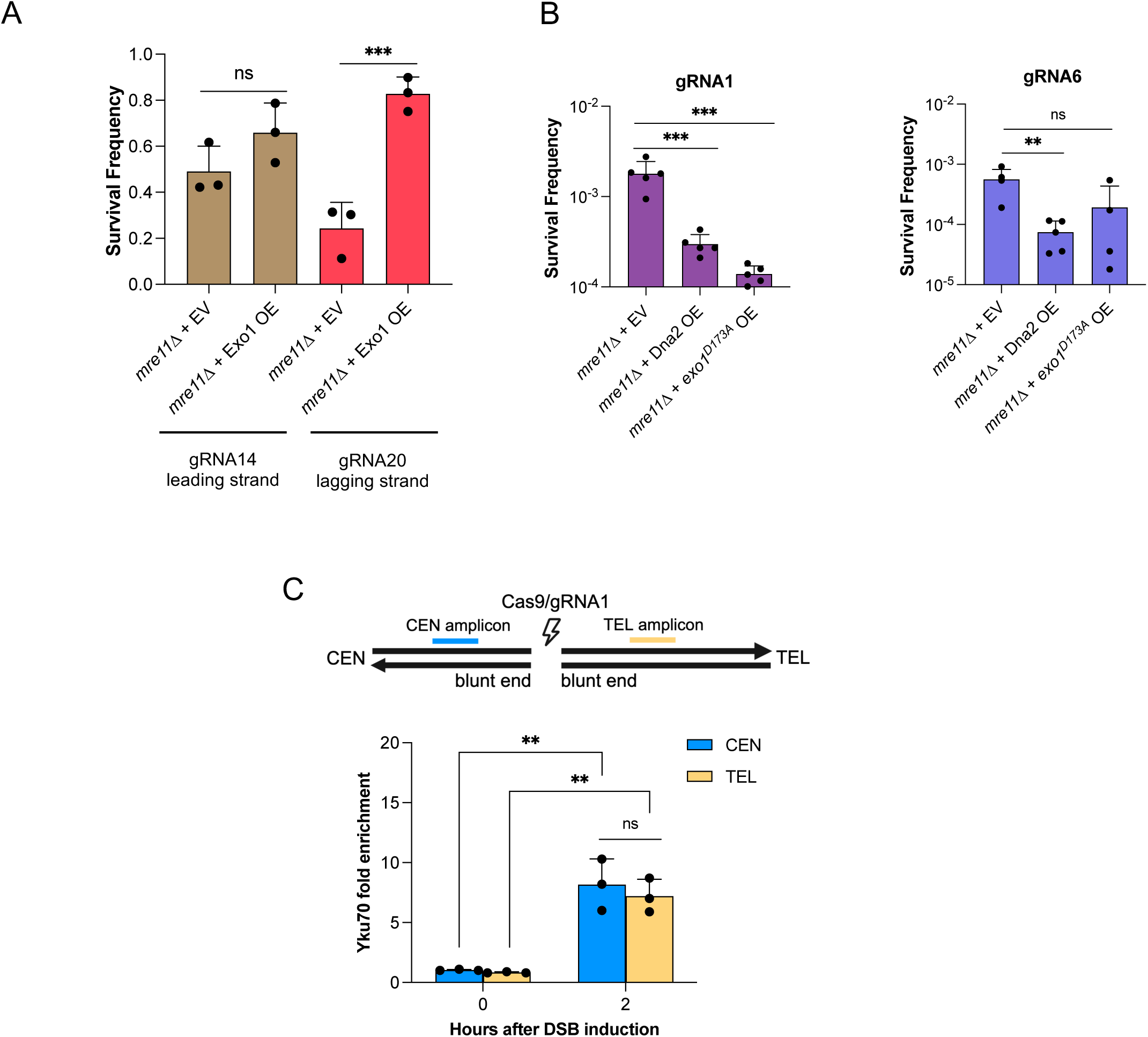
Exo1 overexpression only rescues *mre11*Δ viability to lagging strand DSBs. **A.** Overexpression assays as in Figure 4A with Cas9^D10A^ targeting the gRNA14 and gRNA20 loci. **B.** Survival assays comparing *mre11*Δ + empty vector to *mre11*Δ cells with Dna2 or exo1^D173A^ (nuclease dead) overexpressed in Cas9^D10A^/gRNA1 or gRNA6 targeting conditions. **C.** Yku70 enrichment flanking the gRNA1 locus cut by Cas9. Amplicons to detect Yku70 binding are located 0.1-0.3 kb centromeric or telomeric to the gRNA1 cut site.

**Figure S6.**
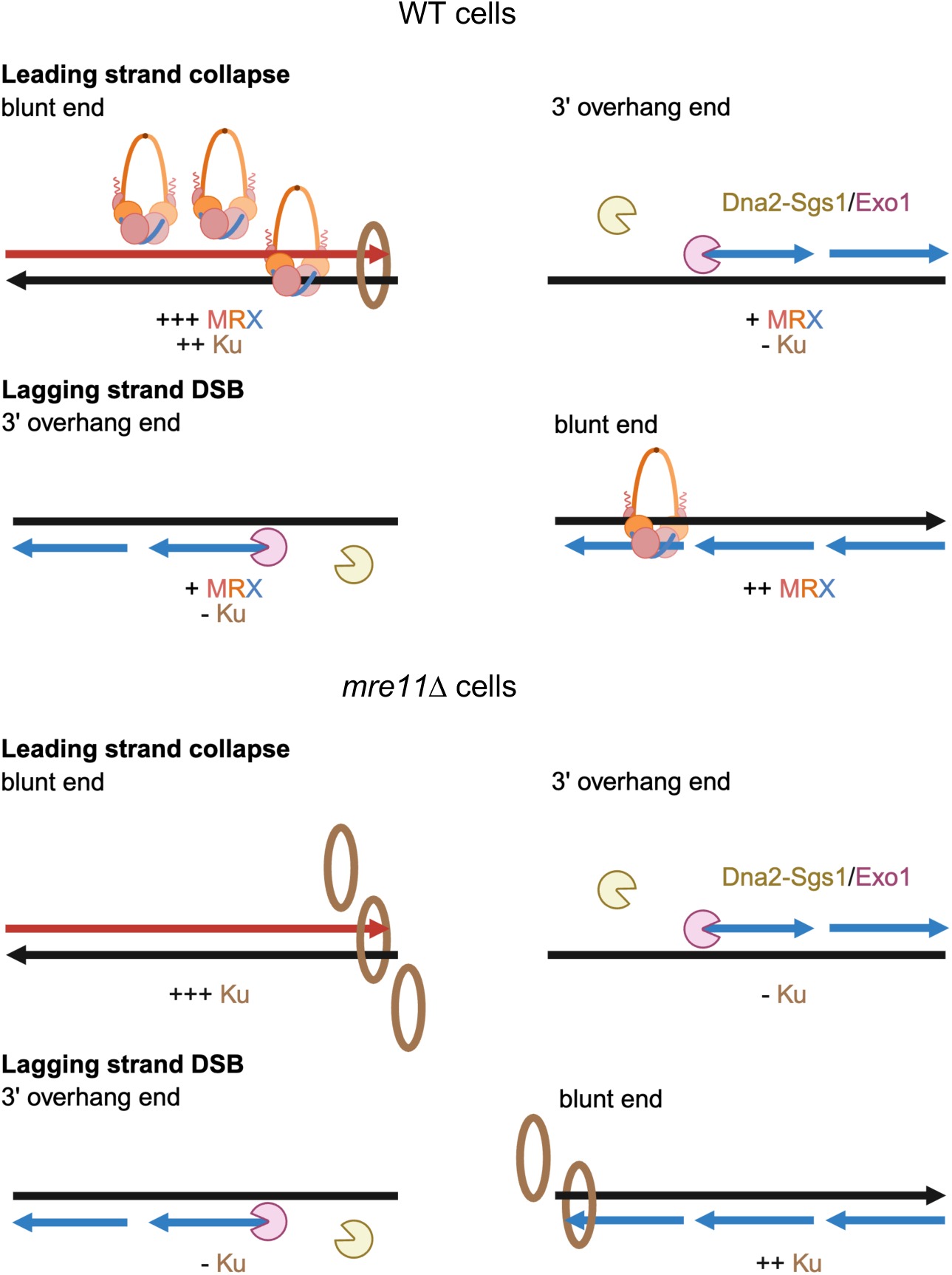
Replication-dependent DSB processing. Model of resection dynamics at both nearly blunt and 3’ overhang DSB ends in WT cells (top) and *mre11*Δ cells (bottom). At ends with 3’ overhangs, Dna2-Sgs1 and Exo1 are still able to initiate long-range resection in the absence of MRX. In contrast, Ku accumulates at nearly blunt ends in the absence of MRX.

**Supplementary Table 1.** Yeast strains.

| Strain # | Genotype* | Source |
| --- | --- | --- |
| LSY0678<br>(W1588-4C) | *MATa* | R. Rothstein |
| LSY1532 | *MATα mre11::TRP1* | Lab stock |
| LSY3397-12C | *MATa rad50::URA3* | Lab stock |
| LSY2992 | *MATα xrs2::KanMX6* | Lab stock |
| LSY3771-1A | *MATα mre11-H125N-NatMX* | Lab stock |
| LSY1091 | *MATa sae2::KanMX6* | Lab stock |
| LSY1996 | *MATa tel1::HphMX4* | Lab stock |
| LSY2374-1 | *MATα exo1::KanMX6* | Lab stock |
| LSY1975 | *MATa sgs1::HphMX4* | Lab stock |
| LSY2401-2B | *MATa exo1::KanMX6 sgs1::HIS3* | Lab stock |
| LSY6182-31D | *MATa mre11-H125N-URA3 exo1::HIS3* | This study |
| LSY6035-7D | *MATα mre11-H125N-NatMX sgs1::HphMX4* | This study |
| LSY4462-1 | *MATa his3::LexO-Cas9<sup>D10A</sup>-3xFLAG-SpHIS5MX*<br>*leu2::ARS607gRNA1-LEU2MX* | Kimble et al.,<br>2025 |
| LSY4469-3 | *MATa his3::LexO-Cas9<sup>D10A</sup>-3xFLAG-SpHIS5MX*<br>*leu2::ARS607gRNA1-LEU2MX mre11::TPR1* | Kimble et al.,<br>2025 |
| LSY6103-9A | *MATa his3::LexO-Cas9<sup>D10A</sup>-3xFLAG-SpHIS5MX*<br>*leu2::ARS607gRNA1-LEU2MX exo1::KanMX sgs1::HphMX* | This study |
| LSY4828-1 | *MATa his3::LexO-Cas9<sup>D10A</sup>-3xFLAG-SpHIS5MX*<br>*leu2::ARS607gRNA6-LEU2MX* | Kimble et al.,<br>2025 |
| LSY4833-2 | *MATa his3::LexO-Cas9<sup>D10A</sup>-3xFLAG-SpHIS5MX*<br>*leu2::ARS607gRNA6-LEU2MX mre11::TRP1* | Kimble et al.,<br>2025 |
| LSY6085-1 | *MATa his3::LexO-Cas9<sup>D10A</sup>-3xFLAG-SpHIS5MX*<br>*leu2::ARS607gRNA6-LEU2MX exo1::KanMX sgs1::HphMX* | This study |
| LSY0785 | *MATa yku70::HIS3* | Lab stock |
| LSY0793 | *MATa mre11::LUE2, yku70::HIS3* | Lab stock |
| LSY5969-1 | *MATa his3::LexO-Cas9<sup>D10A</sup>-3xFLAG-SpHIS5MX*<br>*leu2::ARS607gRNA1-LEU2MX bar1::HphMX4 rad51::KanMX* | This study |
| LSY5589-2 | *MATa his3::LexO-Cas9<sup>D10A</sup>-3xFLAG-SpHIS5MX*<br>*leu2::ARS607gRNA6-LEU2MX rad51::KanMX* | Kimble et al.,<br>2025 |
| LSY6200-1B | *MATα his3::LexO-Cas9<sup>D10A</sup>-SpHIS5MX leu2::ARS607gRNA1-*<br>*LEU2MX, Yku70-6xGly-3xFLAG-HphMX* | This study |
| LSY6201-1C | *MATα his3::LexO-Cas9<sup>D10A</sup>-SpHIS5MX leu2::ARS607gRNA6-*<br>*LEU2MX, Yku70-6xGly-3xFLAG-HphMX* | This study |
| LSY6221-4C | *MATa his3::LexO-Cas9<sup>D10A</sup>-SpHIS5MX leu2::ARS607gRNA1-*<br>*LEU2MX, Yku70-6xGly-3xFLAG-HphMX, mre11::TRP1* | This study |
| LSY6220-16B | *MATα his3::LexO-Cas9<sup>D10A</sup>-SpHIS5MX leu2::ARS607gRNA6-*<br>*LEU2MX, Yku70-6xGly-3xFLAG-HphMX, mre11::TRP1* | This study |
| LSY6181-3A | *MATα his3::LexO-Cas9<sup>D10A</sup>-3xFLAG-SpHIS5MX*<br>*leu2::ARS607gRNA1-LEU2MX mre11-H125N-URA3* | This study |
| LSY6182-2A | *MATa his3::LexO-Cas9<sup>D10A</sup>-3xFLAG-SpHIS5MX*<br>*leu2::ARS607gRNA6-LEU2MX mre11-H125N-URA3* | This study |
| LSY6164-6A | *MATa his3::LexO-Cas9<sup>D10A</sup>-3xFLAG-SpHIS5MX*<br>*leu2::ARS607gRNA1-LEU2MX mre11-H125N-URA3*<br>*sgs1::HphMX* | This study |
| LSY6165-23B | *MATa his3::LexO-Cas9<sup>D10A</sup>-3x-FLAG-SpHIS5MX*<br>*leu2::ARS607gRNA6-LEU2MX mre11-H125N-URA3*<br>*sgs1::HphMX* | This study |
| LSY6181-25B | *MATα his3::LexO-Cas9<sup>D10A</sup>-3x-FLAG-SpHIS5MX*<br>*leu2::ARS607gRNA1-LEU2MX mre11-H125N-URA3 exo1::HIS3* | This study |
| LSY6182-25D | *MATα his3::LexO-Cas9<sup>D10A</sup>-3x-FLAG-SpHIS5MX*<br>*leu2::ARS607gRNA6-LEU2MX mre11-H125N-URA3 exo1::HIS3* | This study |
| LSY4448 | *MATa LexO-Cas9-3x-FLAG-SpHIS5MX leu2::ARS607gRNA1-*<br>*LEU2MX* | Kimble et al.,<br>2025 |
| LSY5768-9B | *MATα his3::LexO-Cas9<sup>D10A</sup>-3x-FLAG-SpHIS5MX*<br>*leu2::ARS607gRNA14-LEU2MX* | Kimble et al.,<br>2025 |
| LSY5771-1A | *MATa his3::LexO-Cas9<sup>D10A</sup>-3x-FLAG-SpHIS5MX*<br>*leu2::ARS607gRNA14-LEU2MX mre11::TRP1* | This study |
| LSY5763-3D | *MATa his3::LexO-Cas9<sup>D10A</sup>-3x-FLAG-SpHIS5MX*<br>*leu2::ARS607gRNA20-LEU2MX* | Kimble et al.,<br>2025 |
| LSY5788-4A | *MATa his3::LexO-Cas9<sup>D10A</sup>-3x-FLAG-SpHIS5MX*<br>*leu2::ARS607gRNA20-LEU2MX mre11::TRP1* | This study |
| LSY4467-2 | *MATa his3::LexO-Cas9<sup>D10A</sup>-3x-FLAG-SpHIS5MX*<br>*leu2::ARS607gRNA1-LEU2MX bar1::HphMX* | Kimble et al.,<br>2025 |
| LSY5869-2A | *MATa his3::LexO-Cas9<sup>D10A</sup>-3x-FLAG-SpHIS5MX*<br>*leu2::ARS607gRNA6-LEU2MX bar1::HphMX* | Kimble et al.,<br>2025 |
| LSY6253-1A | *MATa his3::LexO-Cas9-SpHIS5MX leu2::ARS607gRNA1-*<br>*LEU2MX Yku70-6xGly-3xFLAG-HphMX* | This study |

*\*All strains isogenic to W303, LSY0678 unless otherwise noted*

**Supplementary Table 2-1.**
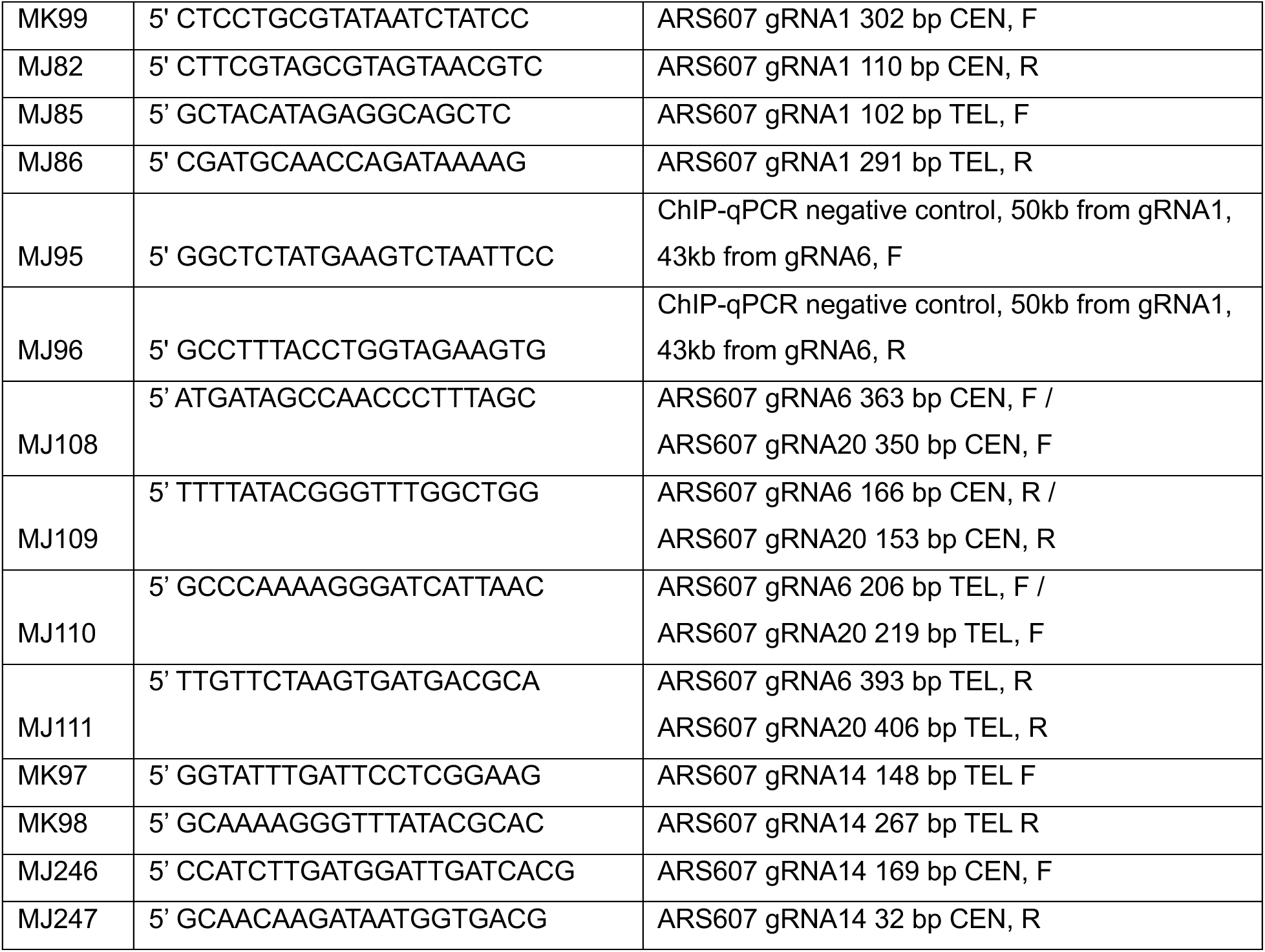
ChIP-qPCR Oligonucleotides.

**Supplementary Table 2-2.**
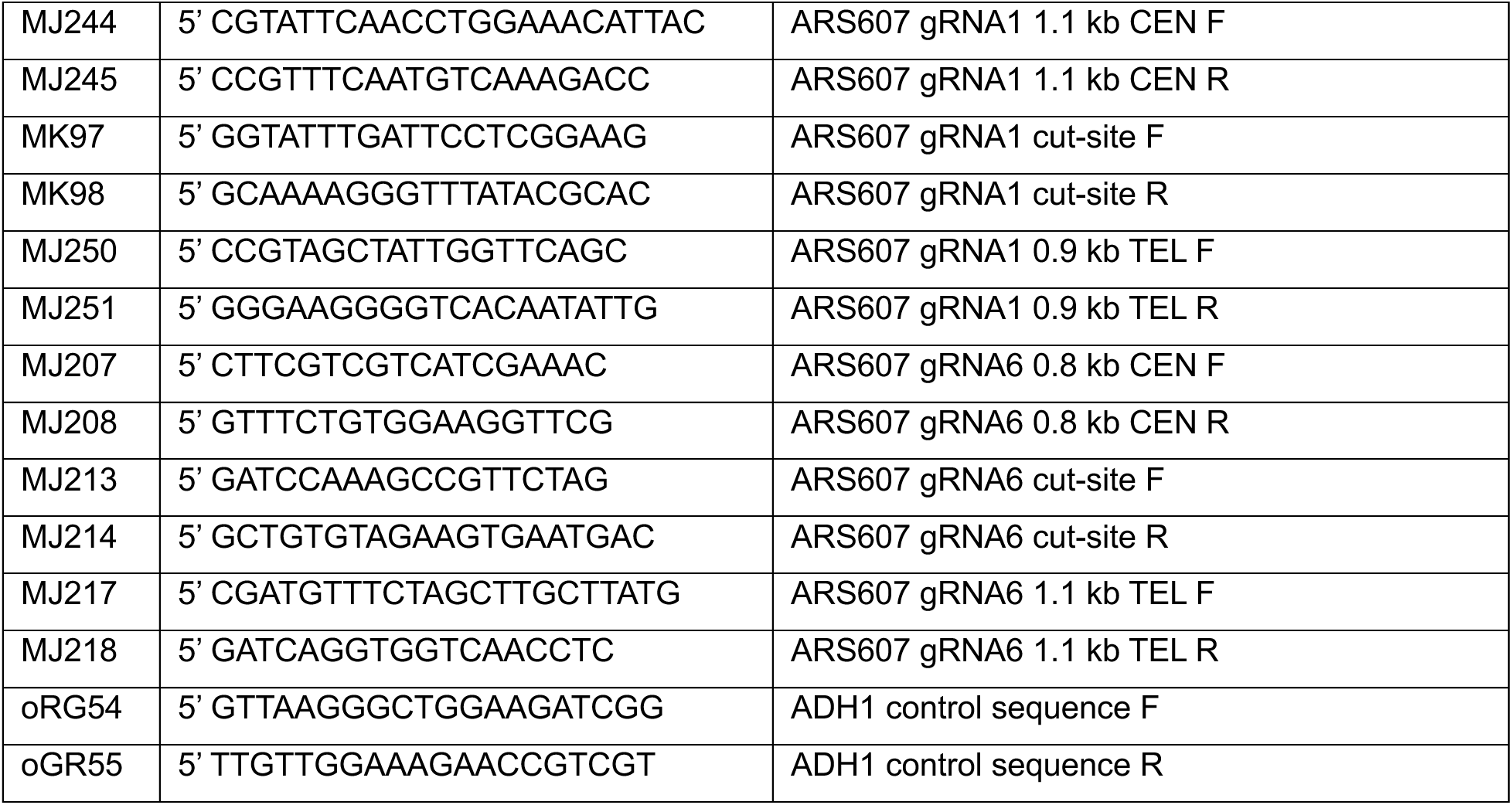
Resection Assay Oligonucleotides.

**Supplementary Table 3.** Plasmids.

| pLS # | Description | I/E<sup>1</sup> | Marker | Source |
| --- | --- | --- | --- | --- |
| 225 | pRS426 | E | URA | Lab stock |
| 14 | EXO1 OE in pRS426 | E | URA | Mimitou and Symington 2010 |
| 667 | DNA2 OE in pRS426 | E | URA | This study |
| 669 | *Exo1*<sup>D173A</sup> OE in pRS426 | E | URA | This study |
| 503 | LexO-CAS9<sup>D10A</sup>-NLS-3xFLAG-T<sub>CYC1</sub> P<sub>ACT1</sub>-LEXA-ERLBD-B112-T<sub>CYC1</sub> in pRG203MX | I | HIS | Kimble et al., 2025 |
| 504 | LexO-CAS9-NLS-3xFLAG-T<sub>CYC1</sub> P<sub>ACT1</sub>-LEXA-ERLBD-B112-T<sub>CYC1</sub> in pRG203MX | I | HIS | Kimble et al., 2025 |
| 517 | LexO-CAS9<sup>D10A</sup>-NLS-3xFLAG-T<sub>CYC1</sub> P<sub>ACT1</sub>-LEXA-ERLBD-B112-T<sub>CYC1</sub> in pRG203MX | I | HIS | Kimble et al., 2025 |
| 627 | LexO-CAS9<sup>D10A</sup>-NLS-3xFLAG-T<sub>ADH1</sub> P<sub>ACT1</sub>-LEXA-ERLBD-B112-T<sub>CYC1</sub> P<sub>tRNA-Tyr</sub>-HDV_ribozyme-gRNA1-T<sub>sNR52</sub> in pRS415 | E | LEU | Kimble et al., 2025 |
| 633 | LexO-CAS9<sup>D10A</sup>-NLS-3xFLAG-T<sub>ADH1</sub> P<sub>ACT1</sub>-LEXA-ERLBD-B112-T<sub>CYC1</sub> P<sub>tRNA-Tyr</sub>-HDV_ribozyme-gRNA6-T<sub>sNR52</sub> in pRS415 | E | LEU | Kimble et al., 2025 |
| 733 | LexO-CAS9<sup>D10A</sup>-NLS-3xFLAG-T<sub>ADH1</sub> P<sub>ACT1</sub>-LEXA-ERLBD-B112-T<sub>CYC1</sub> P<sub>tRNA-Tyr</sub>-HDV_ribozyme-gRNA14-T<sub>sNR52</sub> in pRS415 | E | LEU | Kimble et al., 2025 |
| 735 | LexO-CAS9<sup>D10A</sup>-NLS-3xFLAG-T<sub>ADH1</sub> P<sub>ACT1</sub>-LEXA-ERLBD-B112-T<sub>CYC1</sub> P<sub>tRNA-Tyr</sub>-HDV_ribozyme-gRNA20-T<sub>sNR52</sub> in pRS415 | E | LEU | Kimble et al., 2025 |
| 505 | P<sub>tRNA-Tyr</sub>-HDV_ribozyme-sgRNA-T<sub>sNR52</sub>, with Zral-Xbal gRNA cloning sites in pRG205MX | I | LEU | A. Al-Zain |
| 499 | pLS505 w/ ARS607_gRNA1 | I | LEU | Kimble et al., 2025 |
| 588 | pLS505 w/ ARS607_gRNA6 | I | LEU | Kimble et al., 2025 |
| 708 | pLS505 w/ ARS607_gRNA14 | I | LEU | Kimble et al., 2025 |
| 709 | pLS505 w/ ARS607_gRNA20 | I | LEU | Kimble et al., 2025 |
| 755 | LexO-CAS9<sup>D10A</sup>-NLS-T<sub>CYC1</sub> P<sub>ACT1</sub>-LEXA-ERLBD-B112-T<sub>CYC1</sub> in pRG203MX | I | HIS | This study |
| 758 | LexO-CAS9-NLS-T<sub>CYC1</sub> P<sub>ACT1</sub>-LEXA-ERLBD-B112-T<sub>CYC1</sub> in pRG203MX | I | HIS | This study |

<sup>1</sup> Integrating/Episomal

